# High-order Michaelis-Menten equations allow inference of hidden kinetic parameters in enzyme catalysis

**DOI:** 10.1101/2024.06.12.598609

**Authors:** Divya Singh, Tal Robin, Michael Urbakh, Shlomi Reuveni

## Abstract

Single-molecule measurements provide a platform for investigating the dynamical properties of enzymatic reactions. To this end, the single-molecule Michaelis-Menten equation was instrumental as it asserts that the first moment of the enzymatic turnover time depends linearly on the reciprocal of the substrate concentration. This, in turn, provides robust and convenient means to determine the maximal turnover rate and the Michaelis-Menten constant. Yet, the information provided by these parameters is incomplete and does not allow full characterization of enzyme kinetics at the single molecule level. Here we show that the missing kinetic information can be accessed via a set of high-order Michaelis-Menten equations that we derive. These equations capture universal linear relations between the reciprocal of the substrate concentration and distinguished combinations of turnover time moments, essentially generalizing the Michaelis-Menten equation to moments of any order. We demonstrate how key observables such as the lifetime of the enzyme-substrate complex, the rate of substrate-enzyme binding, and the probability of successful product formation, can all be inferred using these high-order Michaelis-Menten equations.

## Introduction

Enzymes are biocatalysts which enable numerous reactions to take place in living organisms. The Michaelis-Menten mechanism describes their underlying dynamics.^1^ A free enzyme reversibly binds a substrate molecule to form an enzyme-substrate complex that can either lead to the formation of a product via an irreversible catalytic pathway or can simply revert back to the free state without forming a product. Based on this mechanism, the Michaelis-Menten equation predicts a linear relation between the reciprocals of the turnover rate and the substrate concentration. From this linear relation, one can deduce two important kinetic parameters: the maximal rate of the enzymatic reaction and the Michaelis-Menten constant, i.e., the substrate concentration at which half the maximal rate is achieved.^2^

Recent technological advancements have now made it possible to follow the stochastic motion and activity of single particles and molecules^3-11^, including individual enzymes^12-19^, over extended periods of time. Specifically, under certain assumptions, the Michaelis-Menten equation was shown to hold at the single-molecule level where it predicts a linear relation between the mean turnover time and the reciprocal of the substrate concertation.^20-25^ The turnover time probability distribution can also be measured by tracking single enzymes.^12,13,21^ Yet, in contrast to the mean reaction time, higher moments often exhibit complex non-monotonic dependencies on the reciprocal of the substrate concertation^20,21,26,27^, which complicates the extraction of kinetic information and the physical interpretation of experimental results.

Some progress can still be made for enzymes whose kinetics is well described by nearest-neighbor models^27^. Expressions for the mean reaction time and the inverse of the squared coefficient of variation have been attained for such enzymes by adopting a Markovian kinetic description in which internal states are linearly connected in a reversible manner.^28-30^ Using these, information on the number of interim states involved in substrate binding, the total number of states present in the catalytic network, and the number of non-substrate binding states, can be extracted by measuring the mean and second moment of the catalysis time distribution as a function of substrate concentration.^27,29,31,32^ Yet, if the conformational dynamics is non-Markovian^14,33-35^, or has parallel branching pathways or non-sequential motifs, this modeling approach fails.^27^ One is then left to wonder how to systematically extract useful information from higher moments of enzymatic turnover times in a general way and without making restricting assumptions on the underlying kinetics.

Addressing this research problem, several studies have made distinctive progress within the renewal approach to Michaelis-Menten enzyme kinetics^36-40^. In this approach, one adopts the classical reaction scheme of Michaelis & Menten, but allows for non-Markovian transitions between three coarse-grained enzymatic states: free enzyme and substrate (*E* + *S*), enzyme-substrate complex (*ES*), free enzyme and product (*E* + *P*).

The coarse-grained description effectively accounts for multiple intermediate kinetic states that are often part of the reaction, even when information about them is partial or completely missing due to experimental limitations. It can be built by retaining the same state space as in the classical Michaelis-Menten model, while replacing the familiar transition rates with generally distributed (non-Markovian) transition times^41^. Thus, by allowing for arbitrary transition time distributions, one can effectively account for hidden degrees of freedom, e.g., different conformational states and branching pathways, that are lost in the coarse-graining procedure.

A key insight coming from the renewal approach is that the Michaelis-Menten equation is universal.^35^ Namely, regardless of the details of the underlying distributions of binding, unbinding, and catalysis times, the mean turnover time always shows a linear dependence on the reciprocal of the substrate concentration. Yet, extensions of this fundamental result to higher moments of the turnover time have so far remained elusive.

Generalizing the Michaelis-Menten equation to moments of any order will open the door to systematic analysis of turnover time data, thus allowing for characterization of enzyme catalysis beyond the classical, yet incomplete, description of the maximal turnover rate and the Michaelis-Menten constant. Here, we do so by leveraging the renewal approach to identify unique moment combinations that exhibit universal linear dependencies on the reciprocal of the substrate concentration. We call these relations: high-order Michaelis-Menten equations and demonstrate how they can be used to extract kinetic information previously deemed inaccessible, e.g., the binding rate, the mean and variance of the time spent in the bound enzymatic state, and the probability that catalysis occurs before substrate unbinding.

Below, we apply the renewal framework to derive results for the turnover time distribution and continue to show how these yield high-order Michaelis-Menten equations. The benefits coming from these equations, and how to apply them in practice to infer hidden kinetic parameters, are also discussed in detail.

### Renewal approach to enzyme catalysis

Any enzymatic reaction includes three kinetic processes: binding and unbinding of a substrate molecule to and from the enzyme, and catalysis which leads to product formation. These events can be characterized by the random times of binding, *T*_*on*_, unbinding, *T*_*off*_, and of catalysis, *T*_*cat*_ (Fig. 1a). Note that at this point we make no assumptions on the distributions of these times, which in turn determine the experimentally observable turnover time: the time it takes a single enzyme to produce a single molecule of product.

**Figure 1:**
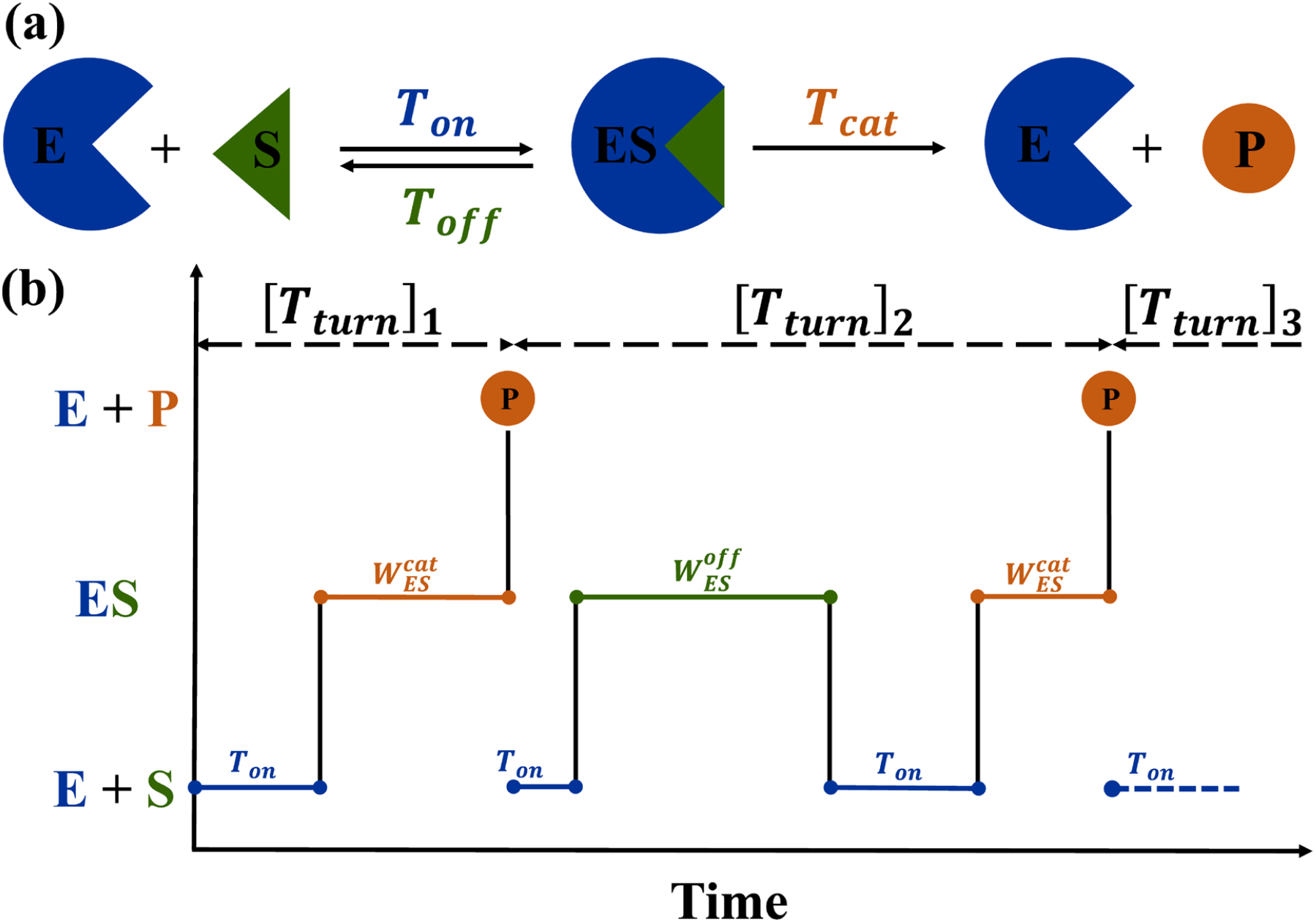
(a) A schematic representation of the Michaelis-Menten model, where transitions are characterized in terms of a general binding time *T*_*on*_, unbinding time *T*_*off*_, and catalysis time *T*_*cat*_. (b) A turnover time trace representing two consecutive product formation events. Following binding, the enzyme is in the *ES* state. If catalysis happens prior to unbinding, a product is formed. In this case, the waiting time in the *ES* state has the statistics of 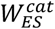 from Eq. (2), i.e., of the catalysis time *T*_*cat*_ *given* that it was smaller than the unbinding time *T*_*off*_. Else, unbinding occurs and the waiting time in the *ES* state has the statistics of 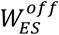 from Eq. (3), i.e., of the unbinding time *T*_*off*_ *given* that it was smaller than the catalysis time *T*_*cat*_. In this paper, we show how to extract hidden information on the statistics of *T*_*on*_, 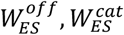, by analyzing the distribution of the turnover time *T*_*turn*_ that is observed in experiments.

The turnover time can be calculated considering two possible routes of enzyme-substrate complex breakup

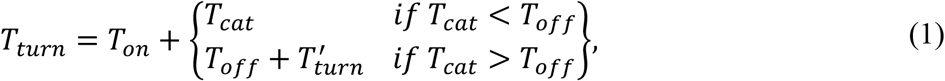

where 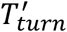 is an identical and independent copy of *T*_*turn*_. Namely, if *T*_*cat*_ is smaller than *T*_*off*_: catalysis follows binding and the enzyme-substrate complex breaks into a free enzyme and product, thus completing the turnover cycle (top row of Eq. (1). Conversely, if *T*_*cat*_ > *T*_*off*_: substrate binding is followed by unbinding and the enzyme-substrate complex breaks into a free enzyme and substrate (bottom row of Eq. (1)). In this case, the turnover cycle starts anew.

To move forward, it is convenient to introduce conditional random variables 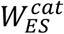 and 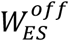 that stand for the waiting times in the enzyme-substrate complex, *provided that either catalysis or unbinding occurs first*, respectively. If catalysis occurs first, the waiting time in the enzyme-substrate complex is given by

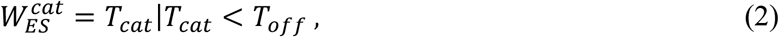

and if unbinding occurs first, the waiting time in the enzyme-substrate complex is given by

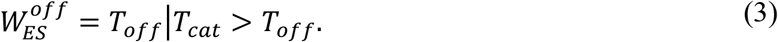

These waiting times have a clear physical meaning which allows for insightful interpretation of the underlying turnover kinetics. In Fig. 1b, we give two possible turnover paths for example. In the first path, binding is directly followed by catalysis and product formation. In the second path, binding is followed by unbinding which restarts the turnover cycle. This is followed by a second binding event leading to catalysis and product formation. We note that single-molecule experiments are designed to track product formation events, but in most cases cannot follow state changes within the catalytic trajectory. Thus, the statistics of *T*_*on*_, 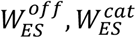 is hidden and the challenge is to infer it from the observed statistics of the turnover times. Next, we develop the theory and practical tools that allow us to do just that.

The probability density functions of the conditional random variables 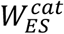 and 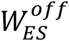 can be written in terms of probability density functions of the catalysis and unbinding times, 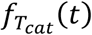 and 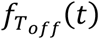, as

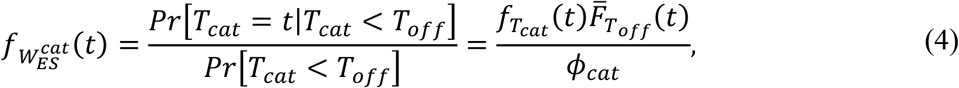

and

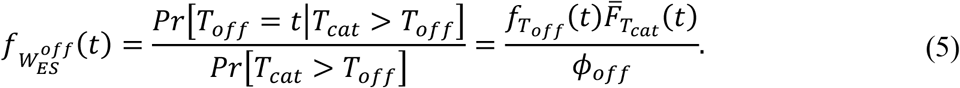

Here 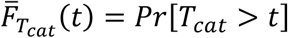 and 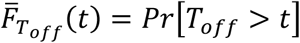 are the survival probability functions of the catalysis and unbinding times respectively, and 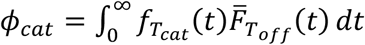 and 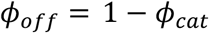 are respectively the probabilities for catalysis to occur before unbinding and vice versa. Naturally, the sum of the latter probabilities is equal to unity. The probability density of the unconditional waiting time at the *ES* state, *W*_*ES*_, is thus given by

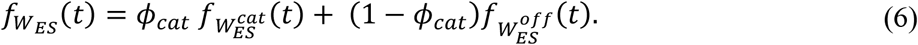

Using the above definitions and assuming that binding times are exponentially distributed with a rate *k*_*on*_ [*S*], the Laplace transform of the turnover time, *T*_*turn*_ in Eq. (1), can be written as (Supplementary Discussion 1)

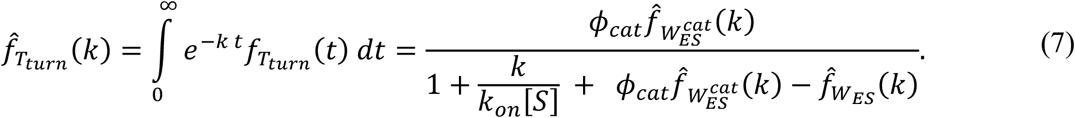

Equation (7) expresses the statistics of turnover times in terms of physically meaningful quantities: the substrate concentration [*SS*], the binding rate *k*_*on*_, the Laplace transforms 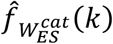 and 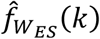 of the conditional and unconditional waiting times in the *ES* state, and the splitting probability *ϕ*_*cat*_.

#### The mean turnover time

The mean turnover time can be computed directly from Eq. (7). It is given by

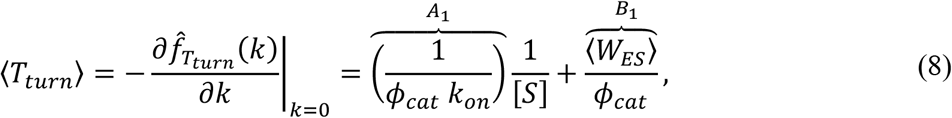

where ⟨*W*_*ES*_⟩ is the mean waiting time in the *ES* state, and all other parameters were already defined. Note that Eq. (8) gives a linear relation between the mean turnover time and the reciprocal substrate concentration. To this end, we define *A*_1_ = 1/(*ϕ*_*cat*_ *k*_*on*_) and *B*_1_ = ⟨*W*_*ES*_ ⟩/ *ϕ*_*cat*_ as the slope and intercept of the corresponding line, respectively.

When binding, unbinding, and catalysis times are all exponentially distributed with rates *k*_*on*_, *k*_*off*_ and *k*_*cat*_, respectively: the mean turnover time in Eq. (8) reduces to the known Michaelis-Menten formula 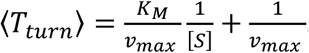. Here *v*_*max*_ = *k*_*cat*_ is the maximal reaction rate and *K*_*M*_ = (*k*_*cat*_ + *k*_*off*_)/ *k*_*on*_ is the Michaelis-Menten constant, which corresponds to the substrate concentration at which half the maximal reaction rate is achieved. This particular case is represented as a blue solid line in Fig. 2a.

**Figure 2:**
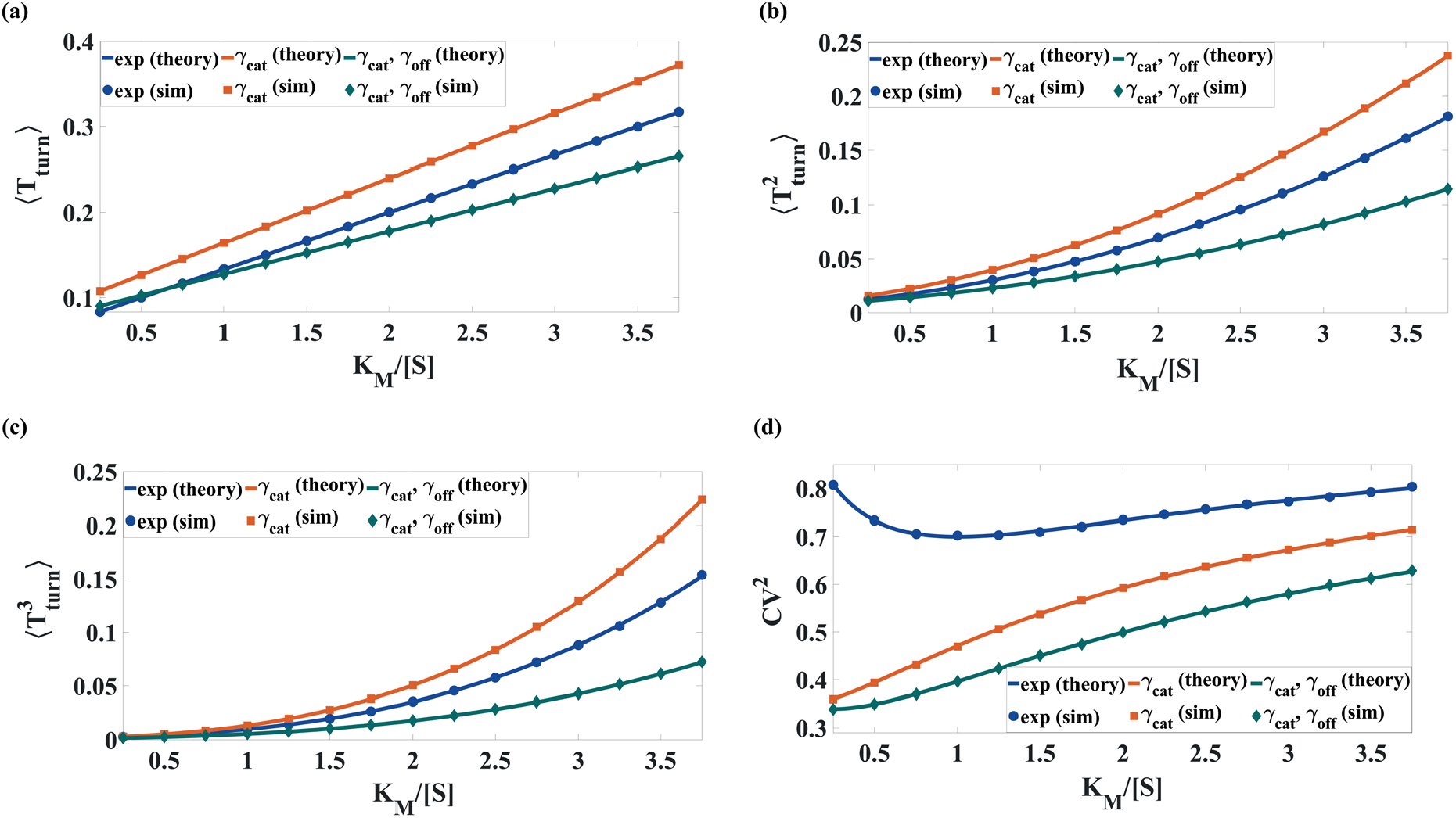
(a) The mean turnover time, ⟨*T*_*turn*_⟩, as a function of the reciprocal substrate concentration (measured in units of the Michaelis constant). We assume a Markovian substrate binding process and consider three different cases for unbinding and catalysis: (i) exponential catalysis and unbinding times (blue), (ii) Gamma distributed catalysis time and exponential unbinding time (orange), and (iii) Gamma distributed catalysis and unbinding times (green). Solid lines come from Eq. (8) which is corroborated by numerical simulations (symbols). In all cases the linear relation predicted by Eq. (8) is shown to hold. (b) Same as (a), but for the second moment ⟨*TT*^2^ ⟩ of the turnover time. (c) Same as (a), but for the third moment 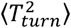 of the turnover time. (d) Same as (a), but for the squared coefficient of variation. It can be appreciated that higher moments, and the squared coefficient of variation, are highly non-linear in the reciprocal substrate concentration.

Along with the classical exponential case, we have also chosen two other enzymatic systems where kinetic processes are non-Markovian. The orange solid line in Fig. 2a represents the case where *T*_*cat*_ follows a Gamma distribution. Similarly, the green solid line represents the case where both *T*_*cat*_ and *T*_*off*_ follow Gamma distributions (see Supplementary Discussion 2 for details). The corresponding symbols represent simulation data points, which are in accord with theory. These results illustrate that the microscopic details of the kinetic transitions do not change the linear dependence on the reciprocal substrate concentration as predicted by Eq. (8).

Summarizing, Eq. (8) extends the basic Michaelis-Menten formula to situations where catalysis and unbinding times come from general distributions. We then have *v*_*max*_ = 1/*B*_1_ = *ϕ*_*cat*_ /⟨*W*_*ES*_ ⟩ and *K*_*M*_ = *A*_1_/*B*_1_ = 1/(*k*_*on*_ ⟨*W*_*ES*_ ⟩). We thus see that the maximal turnover rate is determined by the ratio of the probability, *ϕ*_*cat*_, for a catalytic event to occur before unbinding and the mean waiting time, ⟨*W*_*ES*_⟩, in the *ES* state. Similarly, the Michaelis-Menten constant is determined by the reciprocal of the product between the binding rate and ⟨*W*_*ES*_ ⟩.

#### Michaelis-Menten equations for high turnover time moments

As shown in the previous section, the linear relation between the mean turnover time and the reciprocal substrate concentration (also known as the Lineweaver–Burk plot) allows us to obtain valuable information on enzymatic activity. This is done by fitting a straight line to the data, extracting the slope *A*_1_, and intercept *B*_1_, of Eq. (8); and determining *v*_max_ = 1/*B*_1_ and *K*_*M*_ = *A*_1_/*B*_1_. However, essential microscopic characteristics, such as *k*_*on*_, ⟨*W*_*ES*_⟩ and 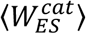 remain unknown. In principle, these parameters can be deduced by analyzing higher moments of turnover time distribution, but this procedure becomes challenging due to the nonlinear dependence of higher moments on 1/[*S*]. These are illustrated in Figs. 2b and 2c for the second and third moments of the turnover times, calculated for various distributions of catalytic and unbinding times. This problem cannot be resolved using commonly considered statistical characteristics of enzymatic activity such as the randomness parameter, Poisson indicator and the Fano factor.^20,26,27,30,42^ As an example, we present in Fig. 2d the dependence of the squared coefficient of variation (randomness parameter), 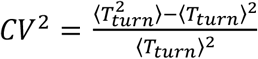, on the reciprocal substrate concentration, which indeed shows a nonlinear behavior for all considered model systems. Notably, even for the classical Michaelis-Menten model, which is characterized by exponential distributions of binding, unbinding and catalytic times, the randomness parameter exhibits a nonmonotonic dependence on 1/[*SS*] (blue line in Fig. 2d).

In order to provide an efficient way to extract microscopic kinetic parameters from single molecule experiments, it is necessary to define combinations of statistical moments that show a linear relationship with the reciprocal substrate concentration. To this end, we take the reciprocal of Eq. (7) and arrive at the following relation

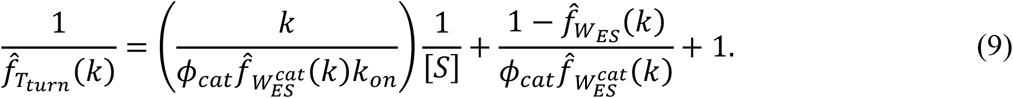

Analogous to the Michaelis-Menten equation reported in Eq. (8), the above equation also shows a linear dependence of 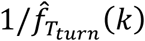 on the reciprocal substrate concentration. Expanding both sides of Eq. (9) in a Taylor series with respect to the Laplace variable *k*, and equating equal order terms, we get (Supplementary Discussion 3):

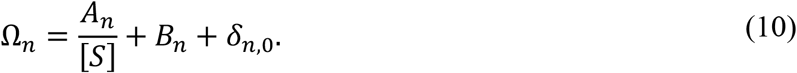

where

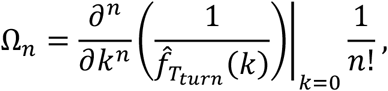

and

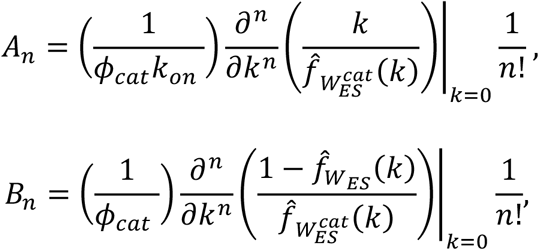

and where *δ*_*n*,0_ is the Kronecker delta.

For *n* = 0 both the left-hand side and right-hand side of Eq. (10) are equal to unity, indicating that all PDFs composing this equation are properly normalized. Also, for *n* = 1, Eq. (10) reduces to Eq. (8) for the first moment. For *n* > 1 the left-hand side of Eq. (10) is a combination of turnover time moments of order {1, …, *n*} whereas its right-hand side is written in terms of kinetic parameters of the enzymatic process (see Table 1 for *n* = 1, 2, and 3 and Supplementary Discussion 4 for further details). Thus, Eq. (10) provides the desired linear relationship between turnover time moments and microscopic parameters of the enzyme. Using an analog of the Lineweaver–Burk plot, the parameters *A*_*n*_ and *B*_*n*_ can be found as the slope and intercept of Ω_*n*_ when plotted vs. 1/[*S*], respectively. Next, we demonstrate these linear relations for the most important cases of *n* = 2 and 3, which include combinations of second and third-order moments. We then proceed to discuss how these relations can be used to infer key parameters of enzymatic catalysis.

**Table 1:** Explicit expressions for Ω_n_, *A*_n_, *B*_n_ from Eq. (10), for *n* = 1,2,3. Note that Ω_n_ is written in terms of *measurable* turnover time moments of order ≤ *n*, and the parameters *A*_n_ and *B*_n_, can be determined as the slope and intercept of Ω_n_ when it is plotted vs. 1/[*S*] respectively. After they are determined, *A*_n_ and *B*_n_ provide two equations for the unknown kinetics parameters for every moment order *n* that is analyzed.

| $n$ | $\Omega_n$ | $A_n$ | $B_n$ |
| --- | --- | --- | --- |
| 1 | $\langle T_{turn} \rangle$ | $\frac{1}{\phi_{cat} k_{on}}$ | $\frac{\langle W_{ES} \rangle}{\phi_{cat}}$ |
| 2 | $\langle T_{turn} \rangle^2 - \frac{\langle T_{turn}^2 \rangle}{2}$ | $\frac{\langle W_{ES}^{cat} \rangle}{\phi_{cat} k_{on}}$ | $\frac{2\langle W_{ES}^{cat} \rangle \langle W_{ES} \rangle - \langle W_{ES}^2 \rangle}{2\phi_{cat}}$ |
| 3 | $\frac{\langle T_{turn}^3 \rangle}{6} - \langle T_{turn} \rangle \langle T_{turn}^2 \rangle + \langle T_{turn} \rangle^3$ | $\frac{(\langle W_{ES}^{cat} \rangle^2 - \frac{\langle (W_{ES}^{cat})^2 \rangle}{2})}{\phi_{cat} k_{on}}$ | $\frac{(\frac{\langle W_{ES}^3 \rangle}{6} - \frac{\langle W_{ES}^{cat} \rangle \langle W_{ES}^2 \rangle}{2} + \langle W_{ES} \rangle \langle W_{ES}^{cat} \rangle^2 - \frac{\langle W_{ES} \rangle \langle (W_{ES}^{cat})^2 \rangle}{2})}{\phi_{cat}}$ |

### Second and third-order Michaelis-Menten equations

Considering second-order terms in Eq. (10), we get the following equation

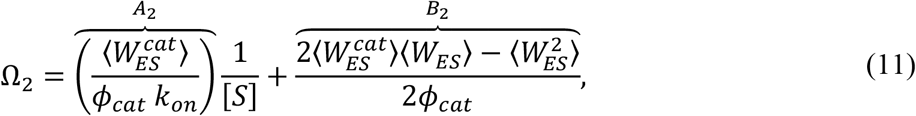

where 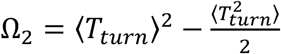 is the specific combination of *n* ≤ 2 turnover time moments, which shows a linear dependence on 1/[*S*] (see Fig. 3). For the classical Michaelis-Menten model, characterized by exponential time distributions, the various PDFs that determine the form of Eq. (10), 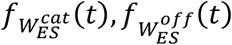 and 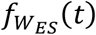, have exactly the same functional form: 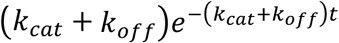 (Supplementary Discussion 5). Thus, in this case 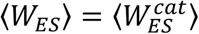 and we have 2 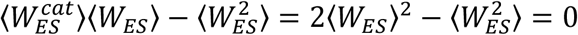, where the last equality follows from basic properties of the exponential distribution. The parameter *B*_2_ which gives the intersection of the line Ω_2_ with the vertical axis, is then equal to zero (blue line in Fig. 3). However, this is not true in general. For non-exponential distributions of catalytic and unbinding times, *B*_2_ can be nonzero (orange and green lines in Fig. 3). Thus, a non-zero value of *B*_2_ can serve as an indicator of the non-classical nature of the enzyme being studied.

**Figure 3:**
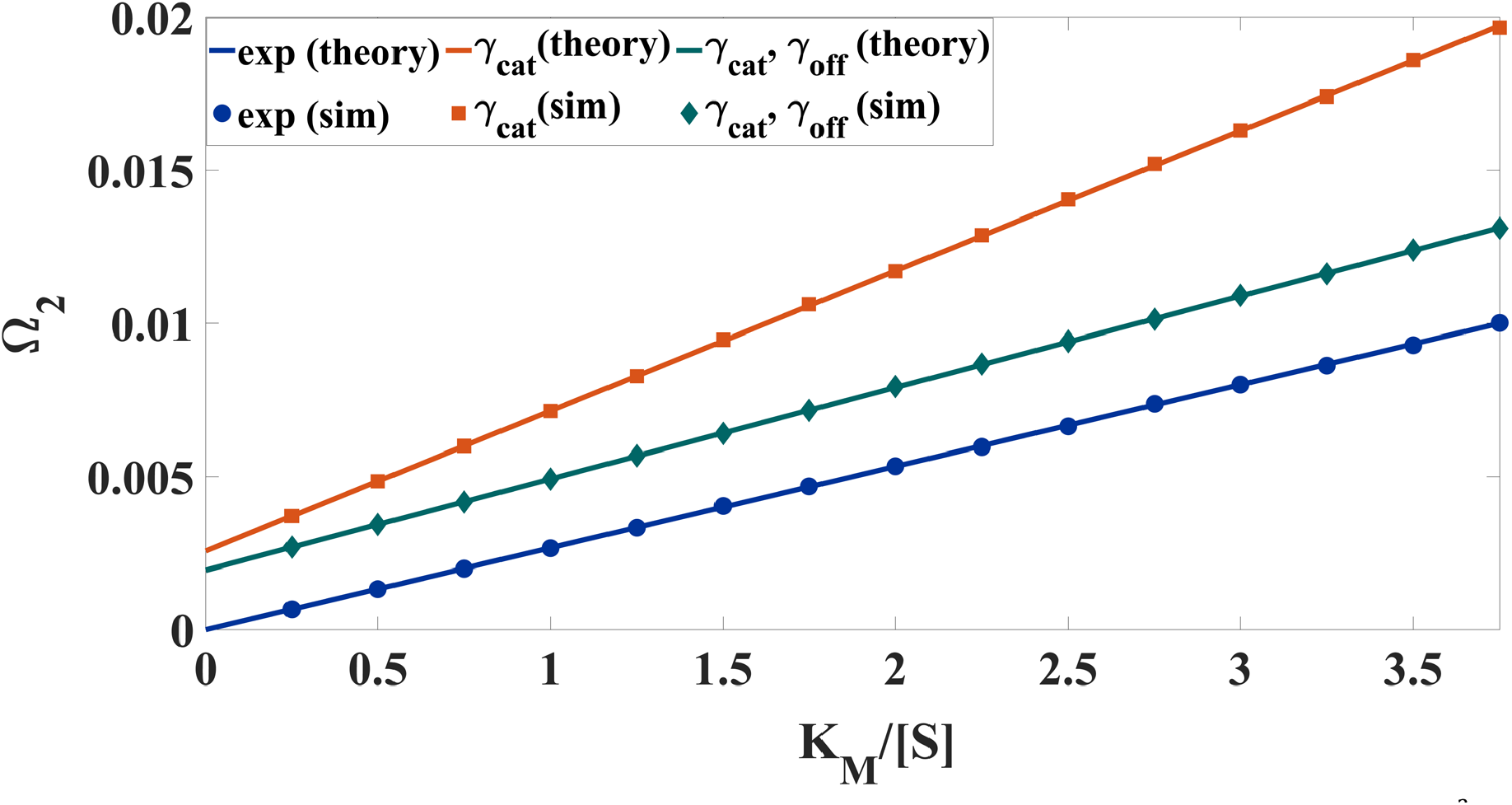
Second order Michaelis-Menten equation. The moments combination, 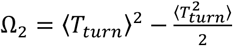, as a function of the reciprocal substrate concentration (measured in units of the Michaelis constant). We assume a Markovian substrate binding process and consider three different cases for unbinding and catalysis: (i) exponential catalysis and unbinding times (blue), (ii) Gamma distributed catalysis time and exponential unbinding time (orange), and (iii) Gamma distributed catalysis and unbinding times (green) [see Supplementary Discussion 2 for details]. Solid lines come from Eq. (11) which is corroborated by numerical simulations (symbols). In all cases the linear relation predicted by Eq. (11) is shown to hold.

Considering third-order terms in Eq. (10) we get the following equation

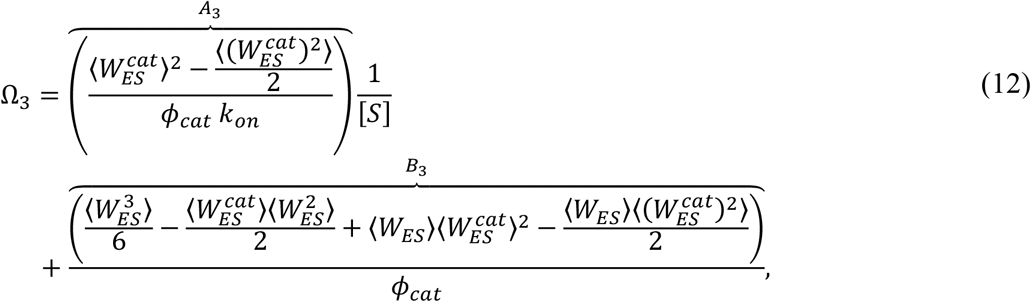

where 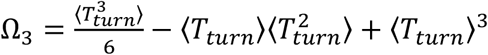 is the specific combination of *n* ≤ 3 turnover time moments, which shows a linear dependence on 1/[*S*] (see Fig. 4). Here too, we see a clear difference between the classical Michaelis-Menten model which is characterized by exponential time distributions, and non-Markovian scenarios. Specifically, in the Markovian case we find that A_3_ = B_3_ = 0 (Supplementary Discussion 6), i.e., the slope and the intercept in Eq. (12) vanish (blue line in Fig. 4). Once again, note that this is not true in general. For non-exponential distributions of catalysis and unbinding times, *A*_3_ and *B*_3_ can be nonzero (orange and green lines in Fig. 4)

**Figure 4:**
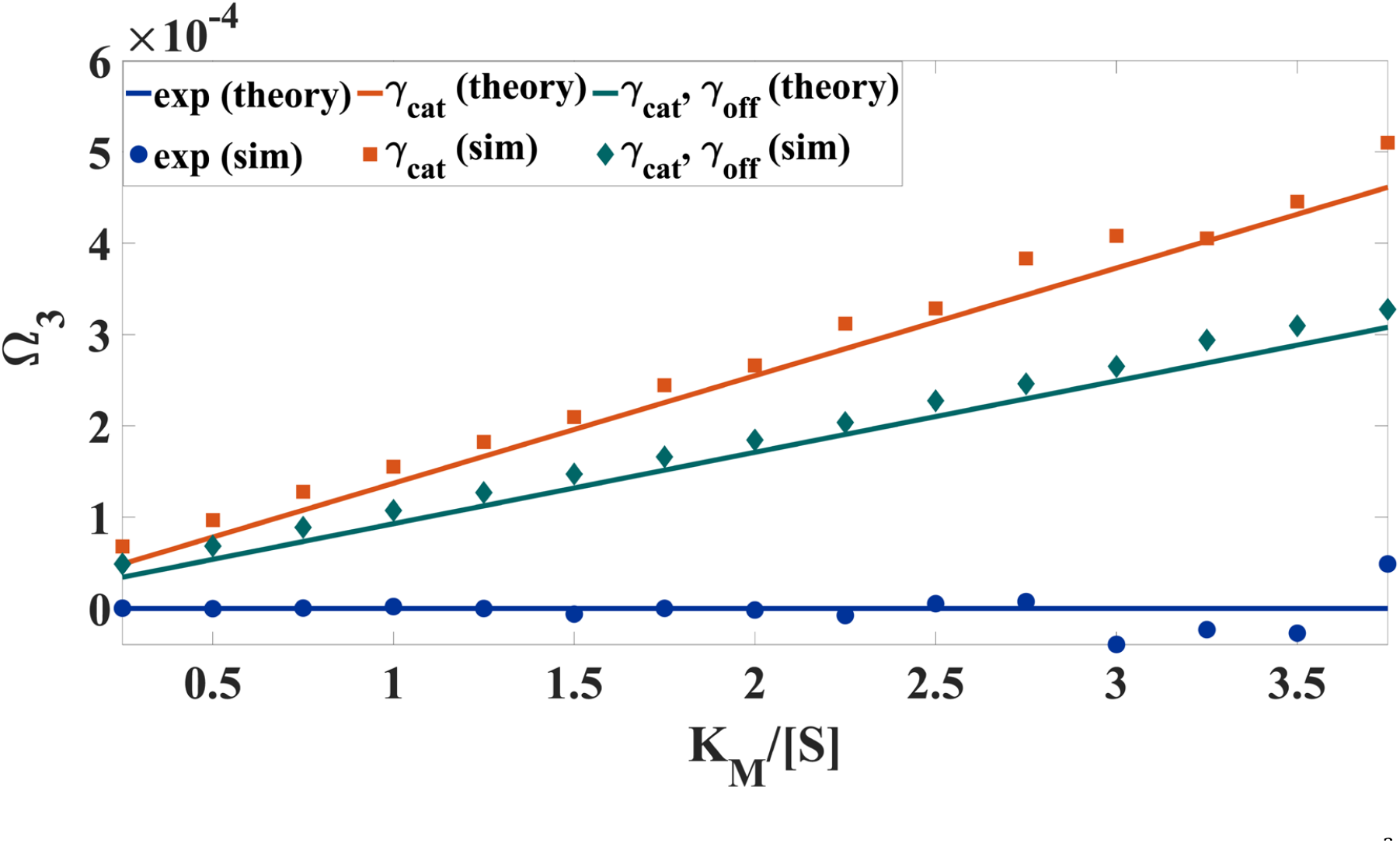
Third order Michaelis-Menten equation. The moments combination, 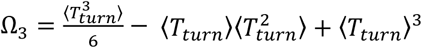, as a function of the reciprocal substrate concentration (measured in units of the Michaelis constant). We assume a Markovian substrate binding process and consider three different cases for unbinding and catalysis: (i) exponential catalysis and unbinding times (blue), (ii) Gamma distributed catalysis time and exponential unbinding time (orange), and (iii) Gamma distributed catalysis and unbinding times (green) [see Supplementary Discussion 2 for details]. Solid lines come from Eq. (12) which is corroborated by numerical simulations (symbols). In all cases the linear relation predicted by Eq. (12) is shown to hold.

### Inferring hidden kinetic parameters from high-order Michaelis-Menten equations

Considering Eqs. (8) and (11) for Ω_1_ and Ω_2_, respectively, we observe that they include four measurable quantities, *A*_1_, *B*_1_, *A*_2_, *B*_2_, and five unknown kinetic parameters, 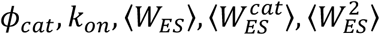, that characterize enzymatic activity (see Table 1). Since an additional equation is missing, these kinetic parameters cannot be uniquely determined only by analyzing the dependence of Ω_1_ and Ω_2_ on the reciprocal substrate concentration. This problem cannot be resolved by considering combinations of higher turnover time moments, Ω_*n*_, with *n* > 2. Indeed, for any *N*, the set of Ω_*n*_, with *n* ≤ *N* allows to obtain 2*N* quantities, *A*_*n*_ and *B*_*n*_, the values of which are determined by 2*N* + 1 unknown kinetic parameters. In particular, plotting Ω_1_, Ω_2_ and Ω_3_ vs. 1/[*S*] one can find six quantities, *A*_*n*_ and *B*_*n*_ for *nn* ≤ 3, which are expressed in terms of seven unknown kinetic parameters, 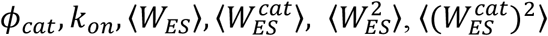 and 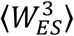 (see Table 1).

From here, one can proceed in multiple ways. First, observe that even if just one of the unknown kinetic parameters can be measured independently, this will provide a closure relation for the set of equations and allow for unique determination of all the other unknown parameters. However, this entails direct measurements of e.g., the splitting probability *ϕ*_*cat*_ or mean time spent at the *ES* state, i.e., ⟨*W*_*ES*_⟩. While such direct measurements may be hard to perform, it is noteworthy that when they are feasible they can be synergistically combined with the framework developed herein to infer hidden kinetic parameters.

Interestingly, missing information can also come from carefully examining other features of the turnover time distribution. In this paper, we focused on moments of the turnover time, but it has recently been shown that short-time behavior also holds valuable kinetic information.^43-45^ Specifically, we have recently shown that the short time behavior of the enzymatic turnover time holds information that allows one to uniquely determine the binding rate *k*_*on*_, provided some mild assumptions hold (Singh, D., Urbakh, M., & Reuveni, S., *Inferring binding rates from enzymatic turnover time statistics*. Manuscript in preparation.) Plugging in the value of the inferred binding rate into the higher-order Michaelis-Menten equations we have developed herein, provides the required closure and allows one to uniquely determine the values of all the other kinetic parameters.

Finally, we show that one can proceed even in the absence of additional information to obtain approximate estimates of the unknown kinetic parameters. In what follows, we show that these approximations are exact for Markovian enzymes, and then proceed to investigate the role of non-Markovianity. For this purpose, using Table 1, it is useful to present the desired quantities, such as the binding rate *k*_*on*_, and the conditional probability for catalysis to happen before unbinding *ϕ*_*cat*_, in the following form

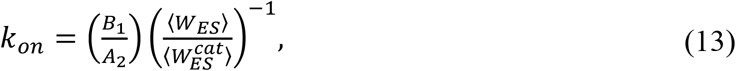

and

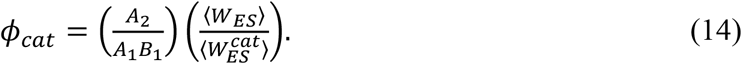

From Table 1, we also see that the mean conditional catalysis time 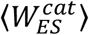 can be calculated using the measured slopes of Ω_1_, and Ω_2_ vs. 1/[*S*] and written as

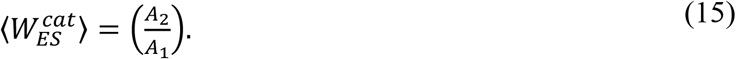

Multiplying both sides of Eq. (15) by 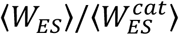, we can write the mean time in the bound enzymatic state as

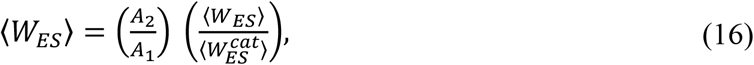

which has a similar form to equations (13) and (14).

Equations (13), (14), and (16) still do not allow us to determine *k*_*on*_ and *ϕ*_*cat*_ and ⟨*W*_*ES*_⟩, since in addition to the measurable quantities, 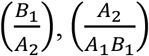 and 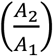 they also include the unknown ratio 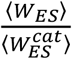. Yet, recall that we have just shown that for classically behaving enzymes, where all the kinetic processes are Markovian, we have 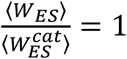. In this case, *k*_*on*_ and *ϕ*_*cat*_ can be directly determined from the slopes (*A*_1_, and *A*_2_) and intercept (*B*_1_) of the Ω_1_ and Ω_2_ lines, as follows

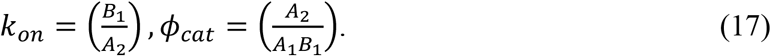

In addition, the rates of unbinding and catalysis can also be inferred from the slopes (*A*_1_, and *A*_2_) and intercept (*B*_1_) as

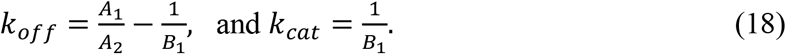

It should be noted that traditional analysis of enzymatic kinetics, which is based only on measurements of the mean turnover times, *cannot* provide this important information.

### Kinetic inference for non-Markovian enzymes

For non-Markovian enzymes, one can *estimate k*_*on*_, *ϕ*_*cat*_, and ⟨*W*_*ES*_⟩ by plugging in 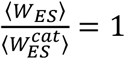in Eqs. (13), (14) and (16). Yet, the quality of this approximation depends on whether the unknown ratio does indeed remain relatively close to unity. In other words, any deviation of 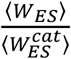 from unity indicates non-Markovian behavior, and the exact value of this ratio provides a measure for the magnitude of the error made when the Markovian approximation is applied. For example, if 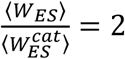, applying the Markovian approximation in Eq. (13) will underestimate the true values of *k*_*on*_ by a factor of two, and similar logic carries to other magnitudes of the error and to *ϕ*_*cat*_ and ⟨*W*_*ES*_⟩ in Eqs. (14) and (16), respectively.

It is thus important to understand in what cases 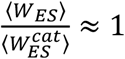 provides a *fair* approximation for non-Markovian enzymes. To answer this question, we now analyze two prevalent, yet distinctly different, types of non-Markovian behavior in enzymes. Namely, non-Markovian behavior that arises due to sequential catalytic steps and non-Markovian behavior that arises due to parallel catalytic pathways. Non-exponential catalysis times are common to both scenarios. Yet, for sequential catalytic steps we have catalysis time distributions that are narrower than the exponential and 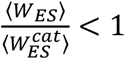, wheras for parallel catalytic pathways we have catalysis time distributions that are broader than the exponential and 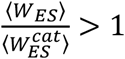.

Let us first consider a model involving two sequential catalytic steps (Fig. 5a). In this model: (i) a free-enzyme, *E*, can reversibly bind a substrate *SS* with rate *k*_*on*_ [*S*], to form an enzyme-substrate complex, *ES*_1_; (ii) the complex *ES*_1_ can then be irreversibly transformed into another complex, *ES*_2_, with rate 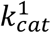 or, alternatively, unbind with rate *k*_*off*_; and finally (iii) the complex *ES*_2_ can be converted by the enzyme to form a product, *P*, with rate 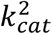 or unbind with rate *k*_*off*_. Note that the kinetic scheme in Fig. 5a can be mapped onto the renewal kinetic scheme in Fig. 1a by coarse graining *ES*_1_ and *ES*_2_ into a single *ES* state and defining *T*_*off*_ and *T*_*cat*_ accordingly (Supplementary Discussion 7). This fact allows us to use the results derived in the previous sections.

**Figure 5:**
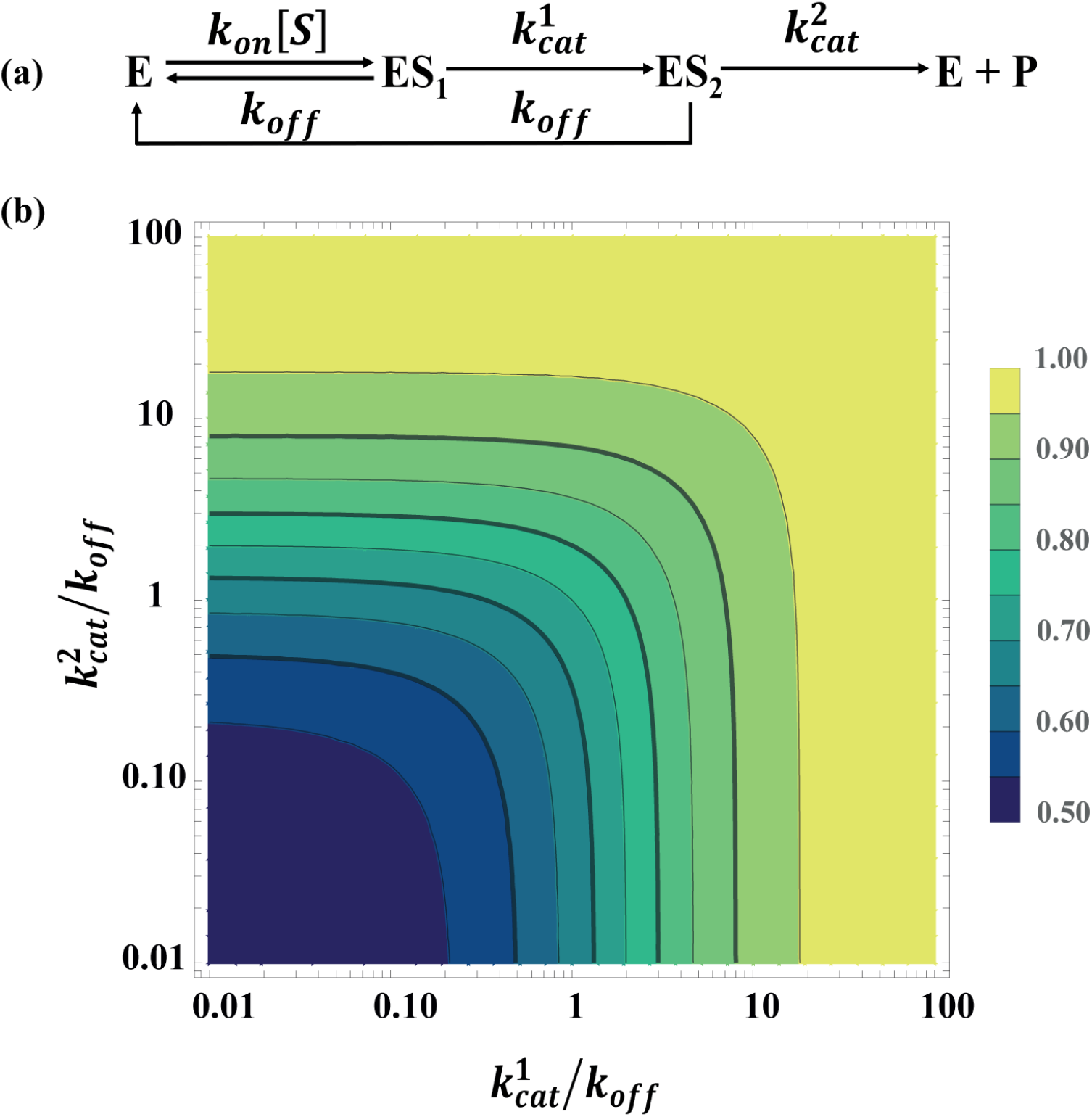
(a) A schematic representation of a model involving two sequential catalytic steps. (b) 2D color map of the error factor 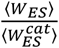 in the space of the normalized catalytic rates 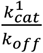 and 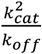.

For the sequential scheme in Fig. 5a, the ratio 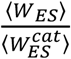 can be calculated analytically, and we find it is given by (Supplementary Discussion 8)

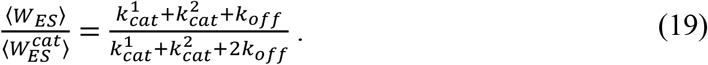

Importantly, Eq. (19) asserts that for any choice of kinetic parameters the ratio lies in the range of 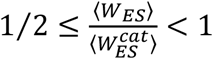, which provides a tight bound on the error coming from the approximation. We visually illustrate this in Figure 5b which presents a 2D color map of 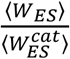, calculated in the space of the normalized catalytic rates 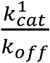 and 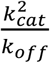. It can be seen that the largest deviations from the classical Markovian behavior, 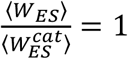, occur when the unbinding rate is much larger than both catalytic rates, 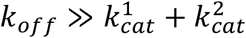.

Summarizing, for enzymes that follow the successive catalytic scheme illustrated in Fig. 5a, the desired kinetic parameters *k*_*on*_ and *ϕ*_*cat*_ can be well estimated by setting 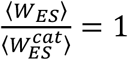 in Eqs. (13) and (14). Also, the mean waiting time in the coarse grained *ES* state can be estimated via Eq. (16) since under the above assumption 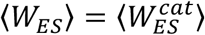. Using these approximations, results in an error factor of no more than two. For example, the Markovian approximation for the binding rate underestimates its true value, but in no more than a factor of two. In the SI, we extend these results to kinetic models with *n* identical sequential steps. We show that the maximal error factor is then given by *n*. Since for real enzymes the value of *n* is rather limited, the Markovian approximation can be applied to sequential enzymes yielding estimates that are well within an order of magnitude of true parameter values (Supplementary Discussion 9).

Next, we consider a reaction scheme involving two parallel catalytic pathways (Fig. 6a). In this model: (i) a free-enzyme, *E*, reversibly binds a substrate, *S*, to form either the enzyme-substrate complex *ES*_1_ or the enzyme-substrate complex *ES*_2_ with rates *pk*_*on*_ [*S*] and (1 − *p*)*k*_*on*_ [*S*], respectively, where *p* is the splitting probability between pathways; (ii) both complexes can then be converted to a product, *P*, with rates 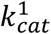 or 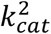, respectively; or unbind with a common rate *k*_*off*_. Unlike the sequential scheme, here the formation of a product can occur from both enzyme-substrate complexes, *ES*_1_ and *ES*_2_. We once again note that the kinetic scheme in Fig. 6a can also be described in the language of Fig. 1a by coarse-graining *ES*_1_ and *ES*_2_ into a single *ES* state and defining *T*_*off*_ and *T*_*cat*_ accordingly (Supplementary Discussion 10). This fact allows us to use the results derived in the previous sections.

**Figure 6:**
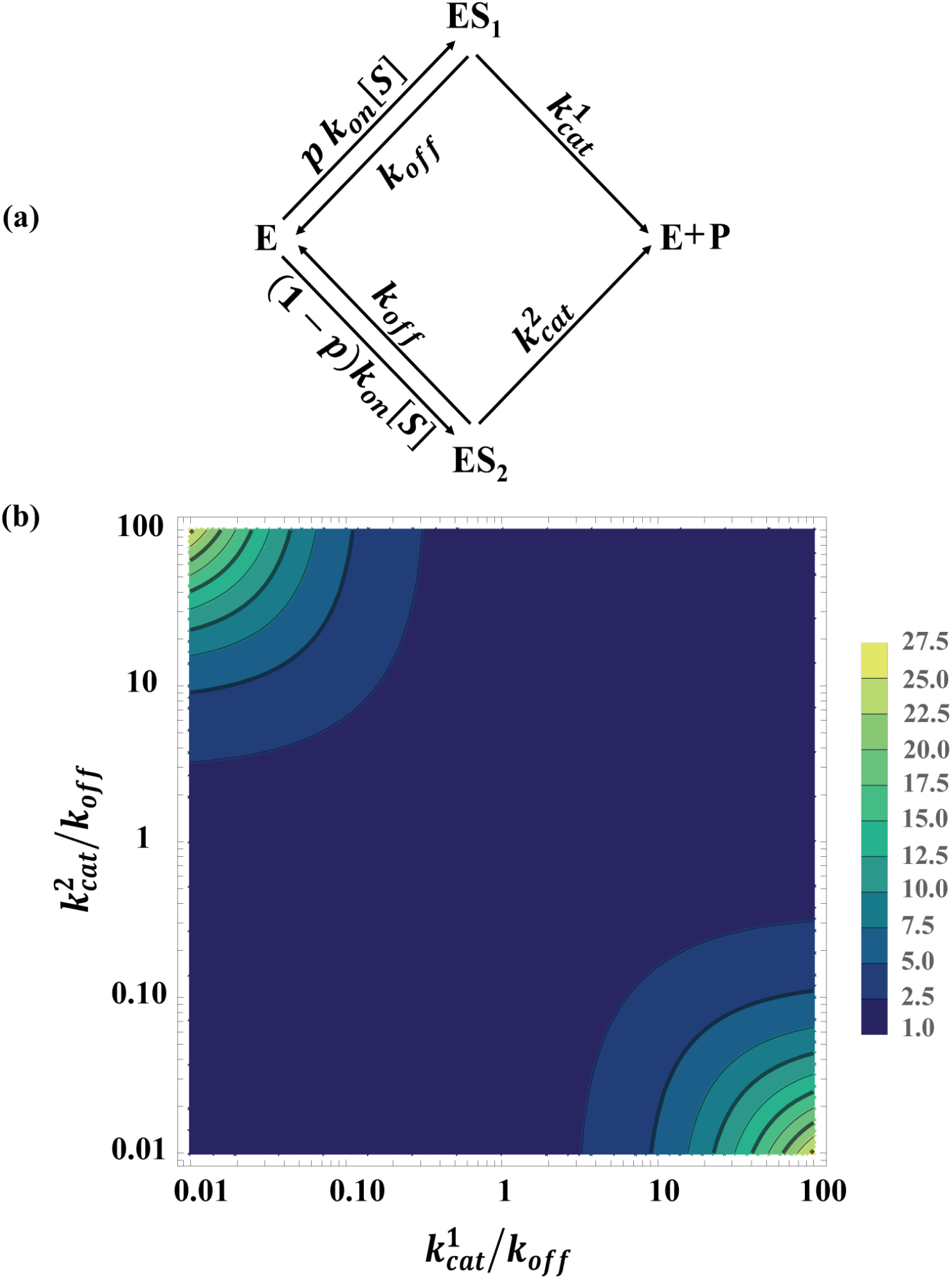
(a) A schematic representation of a model involving two parallel catalytic steps. (b) 2D color map of the error factor 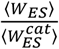 in the space of the normalized catalytic rates 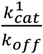 and 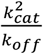, for the splitting probability *p* = 0.5. Note that for 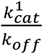 and 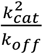 that span four orders of magnitude, the error factor is always <30, and in most cases significantly smaller than this upper bound.

For the parallel kinetic scheme, the ratio 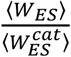 can be calculated analytically, and we find it is given by (Supplementary Discussion 11)

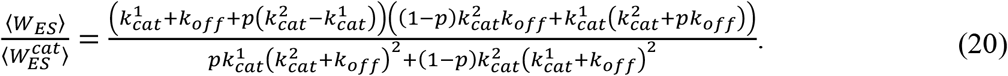

Analysis of this equation asserts that the ratio 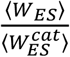 exhibits the strongest deviations from unity for the splitting probability *p* = 0.5, i.e., when the two enzyme-substrate complexes are formed with equal probability. We thus focus on this worst-case scenario.

The corresponding 2D color map of 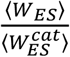, calculated as a function of the normalized catalytic rates 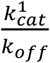and 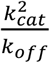 is presented in Fig. 6b. This map clearly demonstrates that in most of the parameter space the ratio 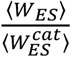 is close to unity, albeit in small corner regions where significant deviations from unity are observed. This happens under extreme conditions where the normalized catalytic rates, 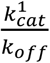 and 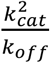, differ from each other by more than three orders of magnitude; and note that in all other cases applying the Markovian approximation in Eqs. (13), (14), and (16) will lead to estimates that are withing an order of magnitude from true values. Similar analysis applies to asymmetric cases where one of the catalytic routes is preferentially chosen over the other (*p* ≠ 0.5), albeit Markovian approximation errors being even smaller (Supplementary Discussion 12). Thus, similar to the sequential scheme, setting 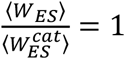 as a closure relation in the parallel kinetic scheme leads to very good approximations for the values of kinetic parameters that are otherwise very hard to estimate.

## Discussion

In this paper, we presented a pioneering approach to enzyme kinetics at the single-molecule level. Building on a renewal approach to enzyme kinetics, we derived a set of high-order Michaelis-Menten equations that reveal universal linear dependencies between unique combinations of turnover time moments and the reciprocal substrate concentration. These relations effectively generalize the single-molecule Michaelis-Menten equation from the mean turnover time to moments of arbitrary order.

We demonstrate the capacity of our approach to infer concealed kinetic parameters, such as the lifetime of the enzyme-substrate complex, the rate of binding, and the probability of successful product formation. With this, we pave the way for systematic characterization of enzyme kinetics from single-molecule data, going well beyond the limited information provided by the all-familiar Michaelis-Menten constant and maximal reaction velocity.

The results we establish apply to Markovian and non-Markovian enzymes alike. For Markovian enzymes, which adhere to memoryless stochastic transitions between states, the framework provides a way to accurately estimate kinetic parameters such as the binding, unbinding and catalysis rates. By leveraging the high-order Michaelis-Menten equations, which capture universal linear dependencies, precise determination of these parameters is achieved.

For non-Markovian enzymes, where kinetic processes exhibit memory effects, our framework is also exact but incomplete. Namely, for inference using our framework to be exact, one of the unknown kinetic parameters, e.g., the binding rate, must be measured or inferred independently. This is not a serious limitation as the binding rate can be inferred based on information held in the short-time tail of the turnover time distribution, provided some mild assumptions hold (Singh, D., Urbakh, M., & Reuveni, S., *Inferring binding rates from enzymatic turnover time statistics*. Manuscript in preparation). Combining such additional information with the results derived herein gives a closed and complete inference framework for non-Markovian enzymes.

Moreover, our framework can be used on non-Markovian enzymes even in lieu of external information on the binding rate or other kinetic parameters. To this end, one assumes that the mean time spent in the enzyme-substrate complex is identical to the (conditional) mean time spent in this complex provided product formation is the direct result of the enzyme-substrate encounter. While this approximation introduces some error, we find that this is typically limited. We demonstrate this by analyzing kinetic models with sequential and parallel product formation pathways. These models display canonical, yet completely orthogonal, types of non-Markovian behavior. Nevertheless, in all cases considered, we show that applying the above assumption provides estimates that are within an order of magnitude from the true value of kinetic parameters that would otherwise remain completely unknown.

In summary, through the derivation of high-order Michaelis-Menten equations, we have shown that traditional analysis methods in enzymology can be extended to unveil central kinetic parameters that have so far remained hidden. We thus offer a systematic and straightforward approach to characterize enzyme kinetics far and beyond the current state of the art.

## Supporting information

Supporting Information for High-order Michaelis-Menten equations allow inference of hidden kinetic parameters in enzyme catalysis

## DATA AVAILABILITY

Data supporting the findings of this manuscript are available from the corresponding author upon reasonable request.

## References

1 Michaelis, L. and Menten, M.L., 1913. Die kinetik der invertinwirkung. Biochem., 49(333-369), p.352.

2 Segel, I.H., 1975. Enzyme kinetics: behavior and analysis of rapid equilibrium and steady state enzyme systems (Vol. 115). New York: Wiley.

3 Moerner, W.E. and Fromm, D.P., 2003. Methods of single-molecule fluorescence spectroscopy and microscopy. Review of Scientific instruments, 74(8), pp.3597–3619.

4 Yang, H., Luo, G., Karnchanaphanurach, P., Louie, T.M., Rech, I., Cova, S., Xun, L. and Xie, X.S., 2003. Protein conformational dynamics probed by single-molecule electron transfer. Science, 302(5643), pp.262–266.

5 Kulzer, F. and Orrit, M., 2004. Single-molecule optics. Annu. Rev. Phys. Chem., 55, pp.585–611.

6 Roy, R., Hohng, S. and Ha, T., 2008. A practical guide to single-molecule FRET. Nature methods, 5(6), pp.507–516.

7 Barkai, E., Garini, Y. and Metzler, R., 2012. Strange kinetics of single molecules in living cells. Physics today, 65(8), pp.29–35.

8 Granik, N., Weiss, L.E., Nehme, E., Levin, M., Chein, M., Perlson, E., Roichman, Y. and Shechtman, Y., 2019. Single-particle diffusion characterization by deep learning. Biophysical journal, 117(2), pp.185–192.

9 Rehfeldt, F. and Weiss, M., 2023. The random walker’s toolbox for analyzing single-particle tracking data. Soft Matter.

10 Scott, S., Weiss, M., Selhuber-Unkel, C., Barooji, Y.F., Sabri, A., Erler, J.T., Metzler, R. and Oddershede, L.B., 2023. Extracting, quantifying, and comparing dynamical and biomechanical properties of living matter through single particle tracking. Physical Chemistry Chemical Physics, 25(3), pp.1513–1537.

11 Schanda, P. and Haran, G., 2024. NMR and Single-Molecule FRET Insights into Fast Protein Motions and Their Relation to Function. Annual Review of Biophysics, 53.

12 Funatsu, T., Harada, Y., Tokunaga, M., Saito, K. and Yanagida, T., 1995. Imaging of single fluorescent molecules and individual ATP turnovers by single myosin molecules in aqueous solution. Nature, 374(6522), pp.555–559.

13 Lu, H.P., Xun, L. and Xie, X.S., 1998. Single-molecule enzymatic dynamics. Science, 282(5395), pp.1877–1882.

14 Flomenbom, O., Velonia, K., Loos, D., Masuo, S., Cotlet, M., Engelborghs, Y., Hofkens, J., Rowan, A.E., Nolte, R.J., Van der Auweraer, M. and de Schryver, F.C., 2005. Stretched exponential decay and correlations in the catalytic activity of fluctuating single lipase molecules. Proceedings of the National Academy of Sciences, 102(7), pp.2368–2372.

15 Wiita, A.P., Perez-Jimenez, R., Walther, K.A., Gräter, F., Berne, B.J., Holmgren, A., Sanchez-Ruiz, J.M. and Fernandez, J.M., 2007. Probing the chemistry of thioredoxin catalysis with force. Nature, 450(7166), pp.124–127.

16 Gumpp, H., Puchner, E.M., Zimmermann, J.L., Gerland, U., Gaub, H.E. and Blank, K., 2009. Triggering enzymatic activity with force. Nano letters, 9(9), pp.3290–3295.

17 Bustamante, C., Cheng, W. and Mejia, Y.X., 2011. Revisiting the central dogma one molecule at a time. Cell, 144(4), pp.480–497.

18 Li, G.W. and Xie, X.S., 2011. Central dogma at the single-molecule level in living cells. Nature, 475(7356), pp.308–315.

19 Grossman-Haham, I., Rosenblum, G., Namani, T. and Hofmann, H., 2018. Slow domain reconfiguration causes power-law kinetics in a two-state enzyme. Proceedings of the National Academy of Sciences, 115(3), pp.513–518.

20 Kou, S.C., Cherayil, B.J., Min, W., English, B.P. and Xie, X.S., 2005. Single-molecule Michaelis-Menten equations. The journal of physical chemistry B, 109(41), pp.19068–19081.

21 English, B.P., Min, W., Van Oijen, A.M., Lee, K.T., Luo, G., Sun, H., Cherayil, B.J., Kou, S.C. and Xie, X.S., 2006. Ever-fluctuating single enzyme molecules: Michaelis-Menten equation revisited. Nature chemical biology, 2(2), pp.87–94.

22 Min, W., Gopich, I.V., English, B.P., Kou, S.C., Xie, X.S. and Szabo, A., 2006. When Does the Michaelis-Menten Equation Hold for Fluctuating Enzymes?. The Journal of Physical Chemistry B, 110(41), pp.20093–20097.

23 Cao, J., 2011. Michaelis-Menten Equation and Detailed Balance in Enzymatic Networks. The Journal of Physical Chemistry B, 115(18), pp.5493–5498.

24 Kolomeisky, A.B., 2011. Michaelis-Menten relations for complex enzymatic networks. The Journal of chemical physics, 134(15).

25 Xie, X.S., 2013. Enzyme kinetics, past and present. Science, 342(6165), pp.1457–1459.

26 Singh, D. and Chaudhury, S., 2017. Statistical properties of fluctuating enzymes with dynamic cooperativity using a first passage time distribution formalism. The Journal of chemical physics, 146(14).

27 Moffitt, J.R. and Bustamante, C., 2014. Extracting signal from noise: kinetic mechanisms from a Michaelis–Menten-like expression for enzymatic fluctuations. The FEBS journal, 281(2), pp.498–517.

28 Chemla, Y.R., Moffitt, J.R. and Bustamante, C., 2008. Exact solutions for kinetic models of macromolecular dynamics. The Journal of Physical Chemistry B, 112(19), pp.6025–6044.

29 Moffitt, J.R., Chemla, Y.R. and Bustamante, C., 2010. Mechanistic constraints from the substrate concentration dependence of enzymatic fluctuations. Proceedings of the National Academy of Sciences, 107(36), pp.15739–15744.

30 Chaudhury, S., Cao, J. and Sinitsyn, N.A., 2013. Universality of Poisson indicator and Fano factor of transport event statistics in ion channels and enzyme kinetics. The Journal of Physical Chemistry B, 117(2), pp.503–509.

31 Moffitt, J.R., Chemla, Y.R., Aathavan, K., Grimes, S., Jardine, P.J., Anderson, D.L. and Bustamante, C., 2009. Intersubunit coordination in a homomeric ring ATPase. Nature, 457(7228), pp.446–450.

32 Yu, J., Moffitt, J., Hetherington, C.L., Bustamante, C. and Oster, G., 2010. Mechanochemistry of a viral DNA packaging motor. Journal of molecular biology, 400(2), pp.186–203.

33 Edman, L. and Rigler, R., 2000. Memory landscapes of single-enzyme molecules. Proceedings of the National Academy of Sciences, 97(15), pp.8266–8271.

34 Berezhkovskii, A.M. and Makarov, D.E., 2018. Single-molecule test for Markovianity of the dynamics along a reaction coordinate. The journal of physical chemistry letters, 9(9), pp.2190–2195.

35 Song, K., Makarov, D.E. and Vouga, E., 2023. Compression algorithms reveal memory effects and static disorder in single-molecule trajectories. Physical Review Research, 5(1), p.L012026.

36 Jung, W., Yang, S. and Sung, J., 2010. Novel chemical kinetics for a single enzyme reaction: relationship between substrate concentration and the second moment of enzyme reaction time. The Journal of Physical Chemistry B, 114(30), pp.9840–9847.

37 Yang, S., Cao, J., Silbey, R.J. and Sung, J., 2011. Quantitative interpretation of the randomness in single enzyme turnover times. Biophysical journal, 101(3), pp.519–524.

38 Reuveni, S., Urbakh, M. and Klafter, J., 2014. The role of substrate unbinding in Michaelis-Menten enzymatic reactions. Proc. Natl. Acad. Sci. 111 (12), 4391.

39 Rotbart, T., Reuveni, S. and Urbakh, M., 2015. Michaelis-Menten reaction scheme as a unified approach towards the optimal restart problem. Physical Review E, 92(6), p.060101.

40 Robin, T., Reuveni, S. and Urbakh, M., 2018. Single-molecule theory of enzymatic inhibition. Nature communications, 9(1), p.779.

41 Klafter, J. and Sokolov, I.M., 2011. First steps in random walks: from tools to applications. OUP Oxford.

42 Singh, D., Punia, B. and Chaudhury, S., 2022. Theoretical Tools to Quantify Stochastic Fluctuations in Single-Molecule Catalysis by Enzymes and Nanoparticles. ACS omega, 7(51), pp.47587–47600.

43 Li, X. and Kolomeisky, A.B., 2013. Mechanisms and topology determination of complex chemical and biological network systems from first-passage theoretical approach. The Journal of chemical physics, 139(14).

44 Li, X., Kolomeisky, A.B. and Valleriani, A., 2014. Pathway structure determination in complex stochastic networks with non-exponential dwell times. The Journal of Chemical Physics, 140(18).

45 Thorneywork, A.L., Gladrow, J., Qing, Y., Rico-Pasto, M., Ritort, F., Bayley, H., Kolomeisky, A.B. and Keyser, U.F., 2020. Direct detection of molecular intermediates from first-passage times. Science advances, 6(18), p.eaaz4642.

