## Supporting Information for High-order Michaelis-Menten equations allow inference of hidden kinetic parameters in enzyme catalysis for "High-order Michaelis-Menten equations allow inference of hidden kinetic parameters in enzyme catalysis"

Contents

- Supplementary Discussion 1: Derivation of Eq. (7)
- Supplementary Discussion 2: Kinetic parameters used in Figs. 2, 3 and 4
- Supplementary Discussion 3: Derivation of Eq. (10)
- Supplementary Discussion 4: Derivation of $A_n$ and $B_n$ in Table I
- Supplementary Discussion 5: Proof that $f_{W_{ES}^{cat}}(t), f_{W_{ES}^{off}}(t)$ and $f_{W_{ES}}(t)$ are identical for Markovian enzymes
- Supplementary Discussion 6: Proof that $B_2$, $A_3$ and $B_3$ in table I are zero for Markovian enzymes
- Supplementary Discussion 7: Mapping the kinetic scheme in Fig. 5a onto the renewal kinetic scheme in Fig. 1a
- Supplementary Discussion 8: Derivation of Eq. (19)
- Supplementary Discussion 9: Error factor for the sequential kinetic scheme with $n$ intermediate states
- Supplementary Discussion 10: Mapping the kinetic scheme in Fig. 6a onto the renewal kinetic scheme in Fig. 1a
- Supplementary Discussion 11: Derivation of Eq. (20)
- Supplementary Discussion 12: Error factor for the parallel kinetic scheme with $p = 0.9$

### 1. Derivation of Eq. (7)

Using Eq. (1) and the conditional waiting times defined in Eqs. (2) and (3) in the main text, we obtain

$$f_{T_{turn}}(t) = \phi_{cat} \int_0^t f_{T_{on}}(t') \, f_{W_{ES}^{cat}}(t-t') \, dt' + (1-\phi_{cat}) \int_0^t g(t') \, f_{T'_{turn}}(t-t') \, dt' \,, \tag{S1}$$

where $g(t') = \int_0^{t'} f_{T_{on}}(t'') f_{W_{ES}^{off}}(t'-t'') \, dt''$. The first integral on the right-hand-side of Eq. (S1) is the convolution of the binding time distribution, $f_{T_{on}}(t)$ with the conditional catalysis time distribution $f_{W_{ES}^{cat}}(t)$. In the second integral, we have $g(t)$ which itself is the convolution of the binding time distribution with the conditional unbinding time distribution $f_{W_{ES}^{off}}(t)$. Since unbinding is followed by a fresh turnover cycle, the function $g(t)$ should also be convolved with the probability distribution, $f_{T'_{turn}}(t)$, of a new turnover time. Laplace transforming Eq. (S1) we obtain

$$\hat{f}_{T_{turn}}(k) = \phi_{cat} \, \hat{f}_{T_{on}}(k) \hat{f}_{W_{ES}^{cat}}(k) + (1-\phi_{cat}) \, \hat{f}_{T_{on}}(k) \hat{f}_{W_{ES}^{off}}(k) \hat{f}_{T'_{turn}}(k), \tag{S2}$$

where we used

$$\hat{g}(k) = \int_0^\infty e^{-kt} \left( \int_0^t f_{T_{on}}(t') f_{W_{ES}^{off}}(t-t') \, dt' \right) dt = \hat{f}_{T_{on}}(k) \hat{f}_{W_{ES}^{off}}(k). \tag{S3}$$

Furthermore, since $T'_{turn}$ is an identical and independent copy of $T_{turn}$ we have $\hat{f}_{T_{turn}}(k) = \hat{f}_{T'_{turn}}(k)$. The Laplace transform of the turnover time distribution can then be written as

$$\hat{f}_{T_{turn}}(k) = \frac{\phi_{cat} \, \hat{f}_{T_{on}}(k) \hat{f}_{W_{ES}^{off}}(k)}{1-(1-\phi_{cat}) \, \hat{f}_{T_{on}}(k) \hat{f}_{W_{ES}^{off}}(k)}. \tag{S4}$$

After some rearrangements, Eq. (S4) can be rewritten as

$$\hat{f}_{T_{turn}}(k) = \frac{\phi_{cat} \hat{f}_{W_{ES}^{cat}}(k)}{\frac{1}{\hat{f}_{T_{on}}(k)} - (1-\phi_{cat}) \hat{f}_{W_{ES}^{off}}(k)}. \tag{S5}$$

To proceed, we Laplace transform Eq. (6) in the main text. This gives

$$\hat{f}_{W_{ES}}(k) = \phi_{cat} \, \hat{f}_{W_{ES}^{cat}}(k) + (1-\phi_{cat}) \hat{f}_{W_{ES}^{off}}(k), \tag{S6}$$

and using this relation to substitute for $(1-\phi_{cat}) \hat{f}_{W_{ES}^{off}}(k)$ in Eq. (S5) gives

$$\hat{f}_{T_{turn}}(k) = \frac{\phi_{cat}\hat{f}_{W_{ES}^{cat}}(k)}{\frac{1}{\hat{f}_{T_{on}}(k)} + \phi_{cat}\hat{f}_{W_{ES}^{cat}}(k) - \hat{f}_{W_{ES}}(k)}. \tag{S7}$$

Eq. (7) in the main text follows by assuming that substrate binding times are exponentially distributed with rate $k_{on}[S]$. Under this assumption, we have

$$\frac{1}{\hat{f}_{T_{on}}(k)} = 1 + \frac{k}{k_{on}[S]}, \tag{S8}$$

and substituting back into Eq. (S7) gives Eq. (7).

### 2. Kinetic parameters used in Figs. 2, 3 and 4

The results presented in Fig. 2 of the main text were obtained considering three different catalytic models: (i) exponential catalysis and unbinding times (blue), (ii) Gamma distributed catalysis times and exponential unbinding times (orange), and (iii) Gamma distributed catalysis and unbinding times (green). The common substrate binding event in all three enzymatic systems is represented by an exponential distribution with a mean binding time of $\langle T_{on}\rangle = (k_{on}[S])^{-1} \equiv (5 * [S])^{-1}$.

In the first case, exponential catalysis and unbinding times, the mean unbinding and catalysis times were taken to be $\langle T_{off}\rangle = \left(k_{off}\right)^{-1} = 1/10$ and $\langle T_{cat}\rangle = (k_{cat})^{-1} = 1/15$, respectively.

In the second case, Gamma distributed catalysis times and exponential unbinding times, the unbinding time distribution was taken to be the same as in the first case, and catalysis times were taken to follow a Gamma distribution of the form, $f_{T_{cat}}(t) = \frac{e^{\frac{-t}{\theta}}t^{k-1}}{\theta^k\Gamma[k]}$, with shape factor $k = 6\frac{2}{3}$ and scale factor $\theta = \frac{1}{100}$. The parameters of the Gamma distribution were chosen such that the mean catalysis time for this system, $\langle T_{cat}\rangle = k\theta = 1/15$, would be identical to the mean catalysis time of the exponential case, $\langle T_{cat}\rangle = \frac{1}{k_{cat}} = 1/15$, thereby allowing for a fair comparison.

In the third case, both the catalytic and unbinding times follow Gamma distributions. The parameters for the catalytic time distribution are the same as in the second case, while the unbinding time distribution has a form, $f_{T_{off}}(t) = \frac{e^{\frac{-t}{\theta}}t^{k-1}}{\theta^k\Gamma[k]}$, with shape factor $k = 10$ and scale factor $\theta = \frac{1}{100}$. Note that these kinetic parameters were intentionally chosen such that the mean unbinding time, $\langle T_{off}\rangle = k\theta = 1/10$, is exactly equal to the inverse of the unbinding rate constant used for the exponential case, $\langle T_{off}\rangle = \frac{1}{k_{off}} = 1/10$.

Finally, we note that the same parameters were used in Figs. 3 and 4. Also, data coming from numerical simulations (symbols) were obtained by taking $10^6$ turnover events at each substrate concentration, for each one of the cases discussed above.

### 3. Derivation of Eq. (10)

Expanding the left-hand side of Eq. (9) from the main text in Taylor series, we get

$$\frac{1}{\hat{f}_{T_{turn}}(k)} = \sum_{n=0}^{\infty} \frac{\partial^n}{\partial k^n} \left( \frac{1}{\hat{f}_{T_{turn}}(k)} \right) \Bigg|_{k=0} \frac{k^n}{n!}. \tag{S9}$$

Now, we differentiate each term on the right-hand side of Eq. (S9) with respect to $k$. This gives

$$\begin{aligned} \frac{1}{\hat{f}_{T_{turn}}(k)} = \frac{1}{\hat{f}_{T_{turn}}(k)}\Bigg|_{k=0} - \frac{1}{\left(\hat{f}_{T_{turn}}(k)\right)^2} \left( \frac{\partial}{\partial k} \hat{f}_{T_{turn}}(k) \right) \Bigg|_{k=0} \frac{k}{1!} \\ + \left\{ - \frac{1}{\left(\hat{f}_{T_{turn}}(k)\right)^2} \left( \frac{\partial^2}{\partial k^2} \hat{f}_{T_{turn}}(k) \right) \Bigg|_{k=0} \right. \\ \left. + \frac{2}{\left(\hat{f}_{T_{turn}}(k)\right)^3} \left( \frac{\partial}{\partial k} \hat{f}_{T_{turn}}(k) \right)^2 \Bigg|_{k=0} \right\} \frac{k^2}{2!} \\ + \left\{ - \frac{1}{\left(\hat{f}_{T_{turn}}(k)\right)^2} \left( \frac{\partial^3}{\partial k^3} \hat{f}_{T_{turn}}(k) \right) \Bigg|_{k=0} \right. \\ + \frac{6}{\left(\hat{f}_{T_{turn}}(k)\right)^3} \left( \frac{\partial}{\partial k} \hat{f}_{T_{turn}}(k) \right) \left( \frac{\partial^2}{\partial k^2} \hat{f}_{T_{turn}}(k) \right) \Bigg|_{k=0} \\ \left. - \frac{6}{\left(\hat{f}_{T_{turn}}(k)\right)^4} \left( \frac{\partial}{\partial k} \hat{f}_{T_{turn}}(k) \right)^3 \Bigg|_{k=0} \right\} \frac{k^3}{3!} + \cdots \end{aligned} \tag{S10}$$

The normalization of the turnover time PDF and known relations between the Laplace transform of a random variable and its moments, give the following equations:

$$\hat{f}_{T_{turn}}(k)\big|_{k=0} = 1 \tag{S11}$$

and

$$\langle T_{turn}^n \rangle = (-1)^n \left( \frac{\partial^n}{\partial k^n} \hat{f}_{T_{turn}}(k) \right) \Bigg|_{k=0}. \tag{S12}$$

Using the above set of equations, we rewrite Eq. (S10) as

$$\begin{aligned} \frac{1}{\hat{f}_{T_{turn}}(k)} = 1 + \langle T_{turn} \rangle k + \left( \langle T_{turn} \rangle^2 - \frac{\langle T_{turn}^2 \rangle}{2} \right) k^2 \\ + \left( \frac{\langle T_{turn}^3 \rangle}{6} - \langle T_{turn} \rangle \langle T_{turn}^2 \rangle + \langle T_{turn} \rangle^3 \right) k^3 + \cdots \end{aligned} \tag{S13}$$

Similarly, the first and second terms on the right-hand side of Eq. (9) of the main text can be written as follows

$$\frac{1}{\phi_{cat}k_{on}}\left(\frac{k}{\hat{f}_{W_{ES}^{cat}}(k)}\right)=\frac{1}{\phi_{cat}k_{on}}\sum_{n=0}^{\infty}\frac{\partial^n}{\partial k^n}\left(\frac{k}{\hat{f}_{W_{ES}^{cat}}(k)}\right)\Bigg|_{k=0}\frac{k^n}{n!} \tag{S14}$$

and

$$\frac{1-\hat{f}_{W_{ES}}(k)}{\phi_{cat}\hat{f}_{W_{ES}^{cat}}(k)}=\frac{1}{\phi_{cat}}\sum_{n=0}^{\infty}\frac{\partial^n}{\partial k^n}\left(\frac{1-\hat{f}_{W_{ES}}(k)}{\hat{f}_{W_{ES}^{cat}}(k)}\right)\Bigg|_{k=0}\frac{k^n}{n!}. \tag{S15}$$

Equation (10) follows from equating equal order $k$ terms on both sides of Eq. (9).

### 4. Derivation of $A_n$ and $B_n$ in Table I

To obtain the expressions in Table I, we differentiate the terms on the right-hand side of Eq. (S14) with respect to $k$. This gives

$$\begin{aligned}\frac{1}{\phi_{cat}k_{on}}\left(\frac{k}{\hat{f}_{W_{ES}^{cat}}(k)}\right)=\frac{1}{\phi_{cat}k_{on}}\Bigg[&\left\{\frac{1}{\hat{f}_{W_{ES}^{cat}}(k)}\Bigg|_{k=0}-\frac{k}{\left(\hat{f}_{W_{ES}^{cat}}(k)\right)^2}\left(\frac{\partial}{\partial k}\hat{f}_{W_{ES}^{cat}}(k)\right)\Bigg|_{k=0}\right\}\frac{k}{1!}+\\ &\left\{\frac{2k}{\left(\hat{f}_{W_{ES}^{cat}}(k)\right)^3}\left(\frac{\partial}{\partial k}\hat{f}_{W_{ES}^{cat}}(k)\right)^2\Bigg|_{k=0}-\frac{2}{\left(\hat{f}_{W_{ES}^{cat}}(k)\right)^2}\left(\frac{\partial}{\partial k}\hat{f}_{W_{ES}^{cat}}(k)\right)\Bigg|_{k=0}-\right.\\ &\left.\frac{k}{\left(\hat{f}_{W_{ES}^{cat}}(k)\right)^2}\left(\frac{\partial^2}{\partial k^2}\hat{f}_{W_{ES}^{cat}}(k)\right)\Bigg|_{k=0}\right\}\frac{k^2}{2!}+\left\{-\frac{6k}{\left(\hat{f}_{W_{ES}^{cat}}(k)\right)^4}\left(\frac{\partial}{\partial k}\hat{f}_{W_{ES}^{cat}}(k)\right)^3\Bigg|_{k=0}+\right.\\ &\frac{6}{\left(\hat{f}_{W_{ES}^{cat}}(k)\right)^3}\left(\frac{\partial}{\partial k}\hat{f}_{W_{ES}^{cat}}(k)\right)^2\Bigg|_{k=0}+\frac{6k}{\left(\hat{f}_{W_{ES}^{cat}}(k)\right)^3}\left(\frac{\partial}{\partial k}\hat{f}_{W_{ES}^{cat}}(k)\right)\left(\frac{\partial^2}{\partial k^2}\hat{f}_{W_{ES}^{cat}}(k)\right)\Bigg|_{k=0}-\\ &\left.\frac{3}{\left(\hat{f}_{W_{ES}^{cat}}(k)\right)^2}\left(\frac{\partial^2}{\partial k^2}\hat{f}_{W_{ES}^{cat}}(k)\right)\Bigg|_{k=0}-\frac{k}{\left(\hat{f}_{W_{ES}^{cat}}(k)\right)^2}\left(\frac{\partial^3}{\partial k^3}\hat{f}_{W_{ES}^{cat}}(k)\right)\Bigg|_{k=0}\right\}\frac{k^3}{3!}+\cdots\Bigg].\end{aligned} \tag{S16}$$

Similarly, differentiating the terms on the right-hand side of Eq. (S15) gives

$$
\begin{aligned}
\frac{1-\hat{f}_{W_{ES}}(k)}{\phi_{cat}\hat{f}_{W_{ES}^{cat}}(k)} = \frac{1}{\phi_{cat}}\Bigg[\Bigg\{ & -\frac{1}{\left(\hat{f}_{W_{ES}^{cat}}(k)\right)}\left(\frac{\partial}{\partial k}\hat{f}_{W_{ES}}(k)\right)\Bigg|_{k=0} \\
& -\frac{\left(1-\hat{f}_{W_{ES}}(k)\right)}{\left(\hat{f}_{W_{ES}^{cat}}(k)\right)^2}\left(\frac{\partial}{\partial k}\hat{f}_{W_{ES}^{cat}}(k)\right)\Bigg|_{k=0}\Bigg\}\frac{k}{1!} \\
& +\Bigg\{\frac{2}{\left(\hat{f}_{W_{ES}^{cat}}(k)\right)^2}\left(\frac{\partial}{\partial k}\hat{f}_{W_{ES}^{cat}}(k)\right)\left(\frac{\partial}{\partial k}\hat{f}_{W_{ES}}(k)\right)\Bigg|_{k=0} \\
& -\frac{1}{\left(\hat{f}_{W_{ES}^{cat}}(k)\right)}\left(\frac{\partial^2}{\partial k^2}\hat{f}_{W_{ES}}(k)\right)\Bigg|_{k=0} + \frac{2\left(1-\hat{f}_{W_{ES}}(k)\right)}{\left(\hat{f}_{W_{ES}^{cat}}(k)\right)^3}\left(\frac{\partial}{\partial k}\hat{f}_{W_{ES}^{cat}}(k)\right)^2\Bigg|_{k=0} \\
& -\frac{\left(1-\hat{f}_{W_{ES}}(k)\right)}{\left(\hat{f}_{W_{ES}^{cat}}(k)\right)^2}\left(\frac{\partial^2}{\partial k^2}\hat{f}_{W_{ES}^{cat}}(k)\right)\Bigg|_{k=0}\Bigg\}\frac{k^2}{2!} \\
& +\Bigg\{\frac{3}{\left(\hat{f}_{W_{ES}^{cat}}(k)\right)^2}\left(\frac{\partial}{\partial k}\hat{f}_{W_{ES}^{cat}}(k)\right)\left(\frac{\partial^2}{\partial k^2}\hat{f}_{W_{ES}}(k)\right)\Bigg|_{k=0} \\
& -\frac{6}{\left(\hat{f}_{W_{ES}^{cat}}(k)\right)^3}\left(\frac{\partial}{\partial k}\hat{f}_{W_{ES}}(k)\right)\left(\frac{\partial}{\partial k}\hat{f}_{W_{ES}^{cat}}(k)\right)^2\Bigg|_{k=0} \\
& +\frac{3}{\left(\hat{f}_{W_{ES}^{cat}}(k)\right)^2}\left(\frac{\partial}{\partial k}\hat{f}_{W_{ES}}(k)\right)\left(\frac{\partial^2}{\partial k^2}\hat{f}_{W_{ES}^{cat}}(k)\right)\Bigg|_{k=0} \\
& -\frac{1}{\left(\hat{f}_{W_{ES}^{cat}}(k)\right)}\left(\frac{\partial^3}{\partial k^3}\hat{f}_{W_{ES}}(k)\right)\Bigg|_{k=0} - \frac{6\left(1-\hat{f}_{W_{ES}}(k)\right)}{\left(\hat{f}_{W_{ES}^{cat}}(k)\right)^4}\left(\frac{\partial}{\partial k}\hat{f}_{W_{ES}^{cat}}(k)\right)^3\Bigg|_{k=0} \\
& +\frac{6\left(1-\hat{f}_{W_{ES}}(k)\right)}{\left(\hat{f}_{W_{ES}^{cat}}(k)\right)^3}\left(\frac{\partial}{\partial k}\hat{f}_{W_{ES}^{cat}}(k)\right)\left(\frac{\partial^2}{\partial k^2}\hat{f}_{W_{ES}^{cat}}(k)\right)\Bigg|_{k=0} \\
& -\frac{\left(1-\hat{f}_{W_{ES}}(k)\right)}{\left(\hat{f}_{W_{ES}^{cat}}(k)\right)^2}\left(\frac{\partial^3}{\partial k^3}\hat{f}_{W_{ES}^{cat}}(k)\right)\Bigg|_{k=0}\Bigg\}\frac{k^3}{3!}+\cdots\Bigg]
\end{aligned}
\tag{S17}
$$

Then, given the normalization of $\hat{f}_{W_{ES}^{cat}}(t)$ and $\hat{f}_{W_{ES}}(t)$ and known relations between the Laplace transform of a random variable and its moments, we get the following equations

$$\hat{f}_{W_{ES}^{cat}}(k)\Big|_{k=0} = 1, \tag{S18}$$

$$\langle (W_{ES}^{cat})^n \rangle = (-1)^n \left( \frac{\partial^n}{\partial k^n} \hat{f}_{W_{ES}^{cat}}(k) \right)\Bigg|_{k=0}, \tag{S19}$$

$$\hat{f}_{W_{ES}}(k)\Big|_{k=0} = 1, \tag{S20}$$

$$\langle W_{ES}^n \rangle = (-1)^n \left( \frac{\partial^n}{\partial k^n} \hat{f}_{W_{ES}}(k) \right)\Bigg|_{k=0}. \tag{S21}$$

Substituting Eqs. (S18)-(S21) into Eqs. (S16) and (S17) we obtain

$$\frac{1}{\phi_{cat} k_{on}} \left( \frac{k}{\hat{f}_{W_{ES}^{cat}}(k)} \right) = \frac{1}{\phi_{cat} k_{on}} \left[ k + \langle W_{ES}^{cat} \rangle k^2 + \left\{ \langle W_{ES}^{cat} \rangle^2 - \frac{\langle (W_{ES}^{cat})^2 \rangle}{2} \right\} k^3 + \cdots \right], \tag{S22}$$

and

$$\frac{1 - \hat{f}_{W_{ES}}(k)}{\phi_{cat} \hat{f}_{W_{ES}^{cat}}(k)} = \frac{1}{\phi_{cat}} \left[ \langle W_{ES} \rangle k + \left\{ \frac{2\langle W_{ES}^{cat} \rangle \langle W_{ES} \rangle - \langle W_{ES}^2 \rangle}{2} \right\} k^2 + \left\{ \frac{\langle W_{ES}^3 \rangle}{6} - \frac{\langle W_{ES}^{cat} \rangle \langle W_{ES}^2 \rangle}{2} + \langle W_{ES} \rangle \langle W_{ES}^{cat} \rangle^2 - \frac{\langle W_{ES} \rangle \langle (W_{ES}^{cat})^2 \rangle}{2} \right\} k^3 + \cdots \right]. \tag{S23}$$

Collecting coefficients corresponding to the same power of $k$ from Eqs. (S22) and (S23) we obtain the analytical expressions for $A_n$ and $B_n$ in Table I. Therefore, the presented mathematical procedure enables Eqs. (S13), (S22), and (S23) that eventually establish Eq. 10 as the general form of Eqs. 8, 11 and 12.

### 5. Proof that $f_{W_{ES}^{cat}}(t), f_{W_{ES}^{off}}(t)$ and $f_{W_{ES}}(t)$ are identical for Markovian enzymes

For the classical Michaelis-Menten reaction, where all the kinetic processes are exponential, the PDFs of substrate unbinding and catalysis times read

$$f_{T_{off}}(t) = k_{off}\, e^{-k_{off} t},$$

and (S24)

$$f_{T_{cat}}(t) = k_{cat}\, e^{-k_{cat} t}.$$

Substituting Eq. (S24) into Eqs. (4) and (5) of the main text gives the following conditional waiting times distributions

$$f_{W_{ES}^{cat}}(t) = \frac{k_{cat}\, e^{-k_{cat}t} e^{-k_{off}t}}{\int_0^\infty k_{cat}\, e^{-k_{cat}t} e^{-k_{off}t}\, dt} = \left(k_{cat} + k_{off}\right) e^{-\left(k_{cat}+k_{off}\right)t}, \tag{S25}$$

and

$$f_{W_{ES}^{off}}(t) = \frac{k_{off}\, e^{-k_{off}t}\, e^{-k_{cat}t}}{\int_0^\infty k_{off}\, e^{-k_{off}t}\, e^{-k_{cat}t}\, dt} = \left(k_{cat} + k_{off}\right) e^{-\left(k_{cat}+k_{off}\right)t}. \tag{S26}$$

Also, in this case the splitting probabilities for catalysis and unbinding are $\phi_{cat} = \frac{k_{cat}}{k_{cat}+k_{off}}$ and $1 - \phi_{cat} = \frac{k_{off}}{k_{cat}+k_{off}}$, respectively. Now, substituting Eqs. (S25) and (S26) into Eq. (6) in the main text, we obtain the PDF of the waiting time distribution function in the $ES$ state. This is given by

$$f_{W_{ES}}(t) = \left(k_{cat} + k_{off}\right) e^{-\left(k_{cat}+k_{off}\right)t}. \tag{S27}$$

Equations (S25), (S26) and (S27) show that within the classical framework, the conditional and unconditional PDFs, $f_{W_{ES}^{cat}}(t)$, $f_{W_{ES}^{off}}(t)$ and $f_{W_{ES}}(t)$, are identical. Hence, we have $\langle (W_{ES}^{cat})^n \rangle = \langle W_{ES}^n \rangle = \langle \left(W_{ES}^{off}\right)^n \rangle$.

### 6. Proof that $B_2$, $A_3$ and $B_3$ in table I are zero for Markovian enzymes

Following Eqs. (S25) and (S27), we can calculate the mean conditional catalytic time and the mean time spent by the enzyme in the bound state, $ES$. Since their functional forms are the same, we get $\langle W_{ES} \rangle = \langle W_{ES}^{cat} \rangle = 1/\left(k_{cat} + k_{off}\right)$. Utilizing basic properties of the exponential distribution we also have $\langle W_{ES}^2 \rangle = \langle (W_{ES}^{cat})^2 \rangle = 2/\left(k_{cat} + k_{off}\right)^2$ and $\langle W_{ES}^3 \rangle = \langle (W_{ES}^{cat})^3 \rangle = 6/\left(k_{cat} + k_{off}\right)^3$. With these expressions at hand, it is easy to see that for Markovian enzymes $B_2 = 2\langle W_{ES}^{cat} \rangle \langle W_{ES} \rangle - \langle W_{ES}^2 \rangle = 0$. Similarly, $A_3 = \langle W_{ES}^{cat} \rangle^2 - \frac{\langle \left(W_{ES}^{cat}\right)^2 \rangle}{2} = 0$ and $B_3 = \frac{\langle W_{ES}^3 \rangle}{6} - \frac{\langle W_{ES}^{cat} \rangle \langle W_{ES}^2 \rangle}{2} + \langle W_{ES} \rangle \langle W_{ES}^{cat} \rangle^2 - \frac{\langle W_{ES} \rangle \langle \left(W_{ES}^{cat}\right)^2 \rangle}{2} = 0$.

### 7. Mapping the kinetic scheme in Fig. 5a onto the renewal kinetic scheme in Fig. 1a

To describe the kinetic scheme in Fig. 5a using the renewal approach to enzyme catalysis, we first need to find the corresponding times for binding, $T_{on}$ , unbinding, $T_{off}$, and catalysis, $T_{cat}$ (Fig. 1a in the main text). By construction, the binding time distribution in Fig. 5a is exponential with rate as $k_{on}[S]$. We thus have

$$f_{T_{on}}(t) = k_{on}[S] e^{-k_{on}[S]t}. \tag{S28}$$

The substrate binding event leads to the first bound state $ES_1$ which is further irreversibly transferred to the second enzyme-substrate complex, $ES_2$. Noting that the substrate unbinds from both these enzyme-substrate complexes at the same rate, $k_{off}$, we have

$$f_{T_{off}}(t) = k_{off} e^{-k_{off} t}, \tag{S29}$$

for the unbinding time distribution. Finally, we note that the product can only be formed from $ES_2$. Thus, the PDF of catalysis times, $f_{T_{cat}}(t)$, is a *convolution* of two exponentially distributed processes with rates $k_{cat}^1$ and $k_{cat}^2$, respectively. We thus have

$$f_{T_{cat}}(t) = \int_0^t \left(k_{cat}^1 e^{-k_{cat}^1 t'}\right) k_{cat}^2 e^{-k_{cat}^2 (t-t')}\, dt' = \frac{\left(e^{-k_{cat}^2 t} - e^{-k_{cat}^1 t}\right) k_{cat}^1 k_{cat}^2}{k_{cat}^1 - k_{cat}^2}. \tag{S30}$$

In this way we have coarse grained the two bound enzymatic states, $ES_1$ and $ES_2$, into a single bound state $ES$, thus transforming from the sequential description of Fig. 5a to the renewal description of Fig. 1a.

**8. Derivation of Eq. (19)**

To derive Eq. (19), we explicitly write the survival probability functions $\left(\bar{F}_X(t) = 1 - \int_0^t f_X(t')\, dt'\right)$ of the unbinding time

$$\bar{F}_{T_{off}}(t) = e^{-k_{off} t}, \tag{S31}$$

and catalysis time

$$\bar{F}_{T_{cat}}(t) = \frac{e^{-k_{cat}^2 t} k_{cat}^1 - e^{-k_{cat}^1 t} k_{cat}^2}{k_{cat}^1 - k_{cat}^2}. \tag{S32}$$

Then, we substitute Eqs. (S31) and (S32) into Eqs. (4), (5), and (6) of the main text to obtain the conditional and unconditional PDFs of the waiting time in the $ES$ state

$$f_{W_{ES}^{cat}}(t) = \frac{e^{-\left(k_{cat}^1 + k_{cat}^2 + k_{off}\right)t}\left(e^{k_{cat}^1 t} - e^{k_{cat}^2 t}\right)\left(k_{cat}^1 + k_{off}\right)\left(k_{cat}^2 + k_{off}\right)}{k_{cat}^1 - k_{cat}^2}, \tag{S33}$$

$$f_{W_{ES}^{off}}(t) = \frac{e^{-\left(k_{cat}^1 + k_{cat}^2 + k_{off}\right)t}\left(e^{k_{cat}^1 t} k_{cat}^1 - e^{k_{cat}^2 t} k_{cat}^2\right)\left(k_{cat}^1 + k_{off}\right)\left(k_{cat}^2 + k_{off}\right)}{\left(k_{cat}^1 - k_{cat}^2\right)\left(k_{cat}^1 + k_{cat}^2 + k_{off}\right)}, \tag{S34}$$

$$f_{W_{ES}}(t) = \frac{e^{-\left(k_{cat}^1 + k_{cat}^2 + k_{off}\right)t}\left(-e^{k_{cat}^2 t} k_{cat}^2\left(k_{cat}^1 + k_{off}\right) + e^{k_{cat}^1 t} k_{cat}^1\left(k_{cat}^2 + k_{off}\right)\right)}{k_{cat}^1 - k_{cat}^2}. \tag{S35}$$

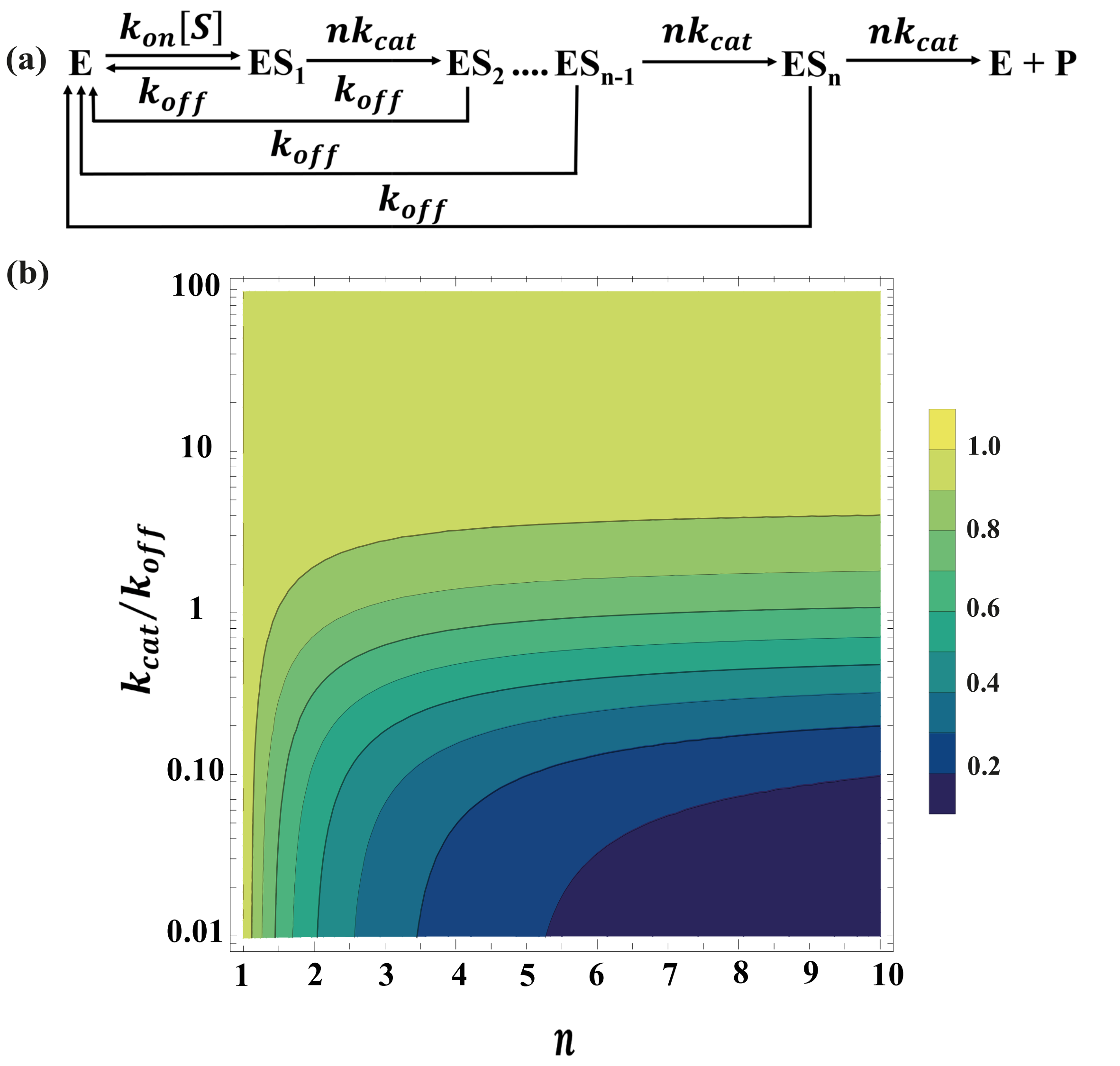


**Figure S1:** (a) A schematic representation of a model involving *n* identical catalytic steps in sequence. (b) 2D color map of the error factor $\frac{\langle W_{ES}\rangle}{\langle W_{ES}^{cat}\rangle}$ calculated in the space of the normalized catalytic rate $\frac{k_{cat}}{k_{off}}$ and the number of catalytic steps $n$ .

The first moments of these distributions can now be computed. These are given by

$$\langle W_{ES}^{cat}\rangle = \frac{1}{k_{cat}^{1} + k_{off}} + \frac{1}{k_{cat}^{2} + k_{off}}, \tag{S36}$$

$$\langle W_{ES}^{off}\rangle = \frac{1}{k_{cat}^{1} + k_{off}} + \frac{1}{k_{cat}^{2} + k_{off}} - \frac{1}{k_{cat}^{1} + k_{cat}^{2} + k_{off}}, \tag{S37}$$

$$\langle W_{ES}\rangle = \frac{k_{cat}^{1} + k_{cat}^{2} + k_{off}}{\left(k_{cat}^{1} + k_{off}\right)\left(k_{cat}^{2} + k_{off}\right)}. \tag{S38}$$

Equation (19) in the main text follows by taking the ratio of $\langle W_{ES}\rangle$ and $\langle W_{ES}^{cat}\rangle$.

### 9. Error factor for the sequential kinetic scheme with $n$ intermediate states

To further evaluate the range of applicability of the Markovian assumption, $\frac{\langle W_{ES}\rangle}{\langle W_{ES}^{cat}\rangle} \approx 1$, we consider here a sequential kinetic scheme with $n$ identical intermediate steps (see Fig. S1a), where each enzyme-substrate complex $ES_i$ can be transformed into the next complex, $ES_{i+1}$, (or in the case of $i = n$ to the product $P$) with rate $nk_{cat}$. Alternatively, at each step unbinding can occur with rate $k_{off}$. Note that rescaling the rate $k_{cat}$ by a factor $n$ is done (without loss of generality) to keep the mean catalytic time equal to $1/k_{cat}$, as in the case of the classical Michaelis-Menten model. This allows for a fair comparison.

To describe the kinetic scheme in Fig. S1a using the renewal approach, we first need to find the corresponding times for binding, $T_{on}$ , unbinding, $T_{off}$, and catalysis, $T_{cat}$. By construction, the binding time distribution in Fig. S1a is exponential with rate $k_{on}[S]$. We thus have $f_{T_{on}}(t) = k_{on}[S]e^{-k_{on}[S]t}$, which is identical to Eq. (28). As all unbinding process follow an exponential distribution with rate $k_{off}$ we have $f_{T_{off}}(t) = k_{off}e^{-k_{off}t}$, which is identical to Eq. (29). Finally, note that from $ES_1$ a chain of irreversible transitions begins to reach the ultimate bound conformer, $ES_n$, which can be transformed into the product. Therefore, the functional form of $f_{T_{cat}}(t)$ is equivalent to the convolution of $n$ sequential steps where each transition occurs with the same rate, $nk_{cat}$. The PDF of catalysis time is thus described by a n-step Erlang distribution function (convolution of n independent and identically distributed exponential steps) which reads

$$f_{T_{cat}}(t) = \frac{(nk_{cat})^n t^{n-1} e^{-nk_{cat}t}}{\Gamma[n]}. \tag{S39}$$

From the above, it follows that the survival probability functions for $f_{T_{cat}}(t)$ and $f_{T_{off}}(t)$ can be written as

$$\bar{F}_{T_{cat}}(t) = \frac{\Gamma[n, nk_{cat}t]}{\Gamma[n]} \tag{S40}$$

and

$$\bar{F}_{T_{off}}(t) = e^{-k_{off}t}. \tag{S41}$$

In Eq. (S40), $\Gamma[n, nk_{cat}t] = \int_{nk_{cat}t}^{\infty} x^{n-1}e^{-x}\,dx$ represents the upper incomplete Gamma function. The algebraic sum of $\Gamma[n, nk_{cat}t]$ and the lower incomplete Gamma function, $\gamma[n, nk_{cat}t] = \int_0^{nk_{cat}t} x^{n-1}e^{-x}\,dx$ gives the complete Gamma function, $\Gamma[n] = \int_0^{\infty} x^{n-1}e^{-x}\,dx$.

Substituting Eqs. (S40) and (S41) into Eqs. (4), (5), and (6) in the main text, we obtain the conditional and unconditional PDFs of the waiting time in the $ES$ state

$$f_{W_{ES}^{cat}}(t) = \frac{\left(nk_{off}\right)^n t^{n-1}\left(\frac{1}{n}+\frac{k_{cat}}{k_{off}}\right)^n e^{-nk_{off}\left(\frac{1}{n}+\frac{k_{cat}}{k_{off}}\right)t}}{\Gamma[n]} \quad \text{(S42)}$$

$$f_{W_{ES}^{off}}(t) = \left(\frac{\Gamma[n, nk_{cat}t]}{\Gamma[n]}\right)\left(\frac{\left(\frac{1}{n}+\frac{k_{cat}}{k_{off}}\right)^n}{\left(\frac{1}{n}+\frac{k_{cat}}{k_{off}}\right)^n - \left(\frac{k_{cat}}{k_{off}}\right)^n}\right) k_{off} e^{-k_{off}t} \quad \text{(S43)}$$

$$f_{W_{ES}}(t) = \frac{\left((nk_{cat})^n t^{n-1} + k_{off}\Gamma[n, nk_{cat}t] e^{nk_{cat}t}\right) e^{-nk_{off}\left(\frac{1}{n}+\frac{k_{cat}}{k_{off}}\right)t}}{\Gamma[n]} \quad \text{(S44)}$$

The corresponding means of these distributions are given by

$$\langle W_{ES}^{cat}\rangle = \frac{1}{k_{off}\left(\frac{1}{n}+\frac{k_{cat}}{k_{off}}\right)} \quad \text{(S45)}$$

$$\langle W_{ES}^{off}\rangle = \frac{1}{k_{off}}\left(1 + \frac{1}{\left(\frac{1}{n}+\frac{k_{cat}}{k_{off}}\right)\left(1-\left(\frac{k_{off}}{k_{cat}}\right)^n\left(\frac{1}{n}+\frac{k_{cat}}{k_{off}}\right)^n\right)}\right), \quad \text{(S46)}$$

and

$$\langle W_{ES}\rangle = \frac{1}{k_{off}}\left(1 - \left(\frac{\frac{k_{cat}}{k_{off}}}{\frac{1}{n}+\frac{k_{cat}}{k_{off}}}\right)^n\right). \quad \text{(S47)}$$

Finally, we obtain the following expression for the ratio $\frac{\langle W_{ES}\rangle}{\langle W_{ES}^{cat}\rangle}$

$$\frac{\langle W_{ES}\rangle}{\langle W_{ES}^{cat}\rangle} = \left(\frac{1}{n}+\frac{k_{cat}}{k_{off}}\right)\left(1 - \left(\frac{\frac{k_{cat}}{k_{off}}}{\frac{1}{n}+\frac{k_{cat}}{k_{off}}}\right)^n\right). \quad \text{(S48)}$$

As expected, in the limit of a single catalytic transition ($n = 1$), this equation gives $\frac{\langle W_{ES}\rangle}{\langle W_{ES}^{cat}\rangle} = 1$. However, for $n > 1$, we find that the error factor in Eq. (S48) is smaller than one but larger than $1/n$ (see Fig. S2). The latter value is obtained in the limit of $k_{cat} \ll k_{off}$ where we have $\langle W_{ES}\rangle \approx 1/k_{off}$ and $\langle W_{ES}^{cat}\rangle \approx n/k_{off}$; and consequently $\langle W_{ES}\rangle/\langle W_{ES}^{cat}\rangle \approx 1/n$.

**10. Mapping the kinetic scheme in Fig. 6a onto the renewal kinetic scheme in Fig. 1**

To describe the kinetic scheme in Fig. 6a using the renewal approach, we again need to find the PDFs of the binding, $T_{on}$ , unbinding, $T_{off}$, and catalysis, $T_{cat}$ times (see Fig.1a in the main text). Here, there are two competitive routes for substrate binding, which occur with probabilities $p$ and $1-p$ to enzyme-substrate complexes, $ES_1$ and $ES_2$, respectively. Therefore, the binding time PDF can be written as $f_{T_{on}}(t) = pk_{on}[S]e^{-k_{on}[S]t} + (1-p)k_{on}[S]e^{-k_{on}[S]t} = k_{on}[S]e^{-k_{on}[S]t}$, which is the same as in the standard Michaelis-

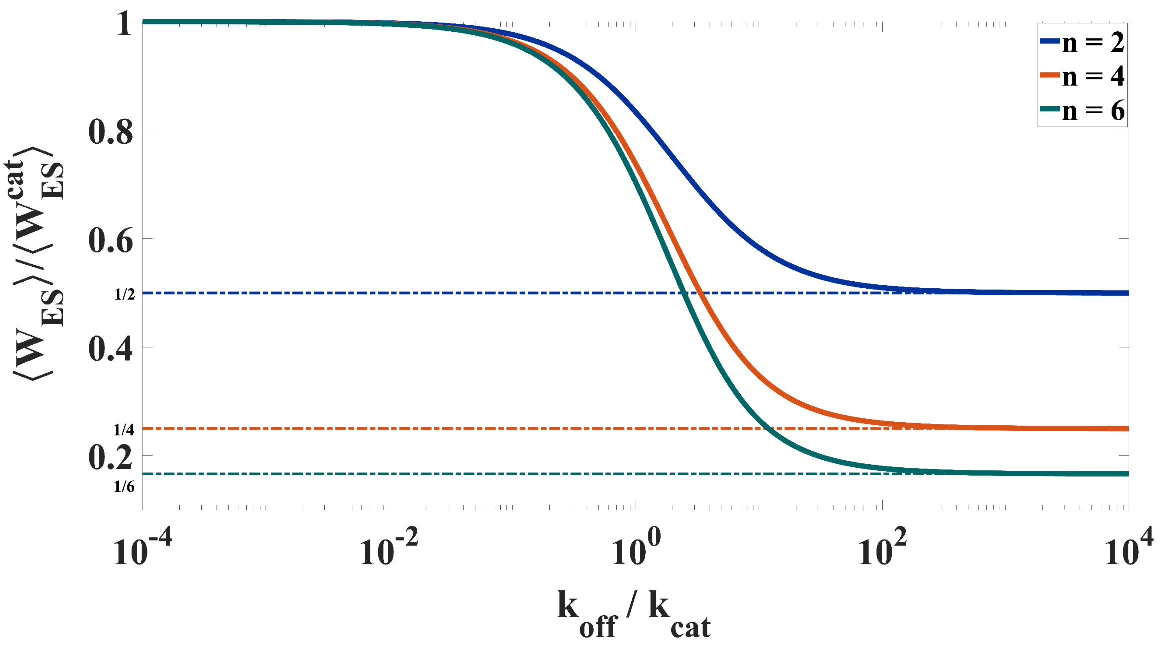


**Figure S2:** The blue, orange and green solid lines show the behavior of $\frac{\langle W_{ES}\rangle}{\langle W_{ES}^{cat}\rangle}$ from Eq. (S48) as a function of $\frac{k_{off}}{k_{cat}}$ for three values of $n$: 2,4, and 6, respectively. Here $n$ is the number of catalytic steps present in the network (see Fig. S1a). We see that the ratio $\frac{\langle W_{ES}\rangle}{\langle W_{ES}^{cat}\rangle}$ decreases with $\frac{k_{off}}{k_{cat}}$, and eventually saturates at $1/n$. The blue, orange, and green dot-dashed lines show the limiting values of $\frac{\langle W_{ES}\rangle}{\langle W_{ES}^{cat}\rangle}$ for the corresponding values of 1/n.

Menten model. Noting that the substrate unbinds from both enzyme-substrate complexes with the same rate, $k_{off}$, we have $f_{T_{off}}(t) = k_{off}e^{-k_{off}t}$ for the unbinding time PDF. Once the $ES_1$ or $ES_2$ complexes are formed, catalysis occurs at the corresponding catalytic rates, namely, $k_{cat}^1$ or $k_{cat}^2$. The catalytic transitions occurring via the two pathways are weighted by the splitting probabilities $p$ and $1-p$, respectively. As a result, the PDF of the catalysis time is given by a double-exponential function.

$$f_{T_{cat}}(t) = p\, k_{cat}^1 e^{-k_{cat}^1 t} + (1-p)k_{cat}^2 e^{-k_{cat}^2 t}. \tag{S49}$$

In this way we have coarse grained the two bound enzymatic states, $ES_1$ and $ES_2$, into a single bound state $ES$, from which product formation occurs with probabilities $p$ at rate $k_{cat}^1$ and with probability $1-p$ at rate $k_{cat}^2$.

### 11. Derivation of Eq. (20)

To derive Eq. (20), we explicitly write the survival probability functions $\left(\bar{F}_X(t) = 1 - \int_0^t f_X(t')\,dt'\right)$ of the catalysis and unbinding times as

$$\bar{F}_{T_{cat}}(t) = pe^{-k_{cat}^1 t} + (1-p)e^{-k_{cat}^2 t} \tag{S50}$$

$$\bar{F}_{T_{off}}(t) = e^{-k_{off}t} \tag{S51}$$

Substituting Eqs. (S50) and (S51) into Eqs. (4), (5) and (6) of the main text, we obtain the following equations

$$f_{W_{ES}^{cat}}(t) = \frac{e^{-(k_{cat}^1+k_{cat}^2+k_{off})t}(k_{cat}^1+k_{off})(k_{cat}^2+k_{off})\left(pk_{cat}^1 e^{k_{cat}^2 t}+(1-p)k_{cat}^2 e^{k_{cat}^1 t}\right)}{pk_{cat}^1(k_{cat}^2+k_{off})+(1-p)k_{cat}^2(k_{cat}^1+k_{off})}, \tag{S52}$$

$$f_{W_{ES}^{off}}(t) = \frac{e^{-(k_{cat}^1+k_{cat}^2+k_{off})t}(k_{cat}^1+k_{off})(k_{cat}^2+k_{off})\left(pe^{k_{cat}^2 t}+(1-p)e^{k_{cat}^1 t}\right)}{p(k_{cat}^2+k_{off})+(1-p)(k_{cat}^1+k_{off})}, \tag{S53}$$

$$f_{W_{ES}}(t) = e^{-(k_{cat}^1+k_{cat}^2+k_{off})t}\left(pe^{k_{cat}^2 t}(k_{cat}^1+k_{off}) + (1-p)e^{k_{cat}^1 t}(k_{cat}^2+k_{off})\right). \tag{S54}$$

The corresponding means of these distributions are given by

$$\langle W_{ES}^{cat}\rangle = \frac{(k_{cat}^1+k_{off})(k_{cat}^2+k_{off})\left(\frac{pk_{cat}^1}{(k_{cat}^1+k_{off})^2}+\frac{(1-p)k_{cat}^2}{(k_{cat}^2+k_{off})^2}\right)}{pk_{cat}^1(k_{cat}^2+k_{off})+(1-p)k_{cat}^2(k_{cat}^1+k_{off})}, \tag{S55}$$

$$\langle W_{ES}^{off}\rangle = \frac{(k_{cat}^1+k_{off})(k_{cat}^2+k_{off})\left(\frac{p}{(k_{cat}^1+k_{off})^2}+\frac{(1-p)}{(k_{cat}^2+k_{off})^2}\right)}{p(k_{cat}^2+k_{off})+(1-p)(k_{cat}^1+k_{off})}, \tag{S56}$$

$$\langle W_{ES}\rangle = \frac{p}{(k_{cat}^1+k_{off})}+\frac{(1-p)}{(k_{cat}^2+k_{off})}. \tag{S57}$$

Using the above expressions, we arrive at Eq. (20) for the ratio of $\langle W_{ES}\rangle$ and $\langle W_{ES}^{cat}\rangle$ that is presented in the main text.

### 12. Error factor for the parallel kinetic scheme with probability $p = 0.9$

As noted in the main text, when the splitting probability is different from 0.5, one catalytic pathway is preferred over the other and deviations of the error factor $\frac{\langle W_{ES}\rangle}{\langle W_{ES}^{cat}\rangle}$ from unity become less significant. We illustrate this in Fig. S3 for a splitting probability $p = 0.9$ which provides a strong bias towards the first catalytic pathway that leads to the $ES_1$ complex. In this case for $\frac{k_{cat}^1}{k_{off}}$ and $\frac{k_{cat}^2}{k_{off}}$ that differ from each other by less than four orders of magnitude, the error factor is $< 10$.

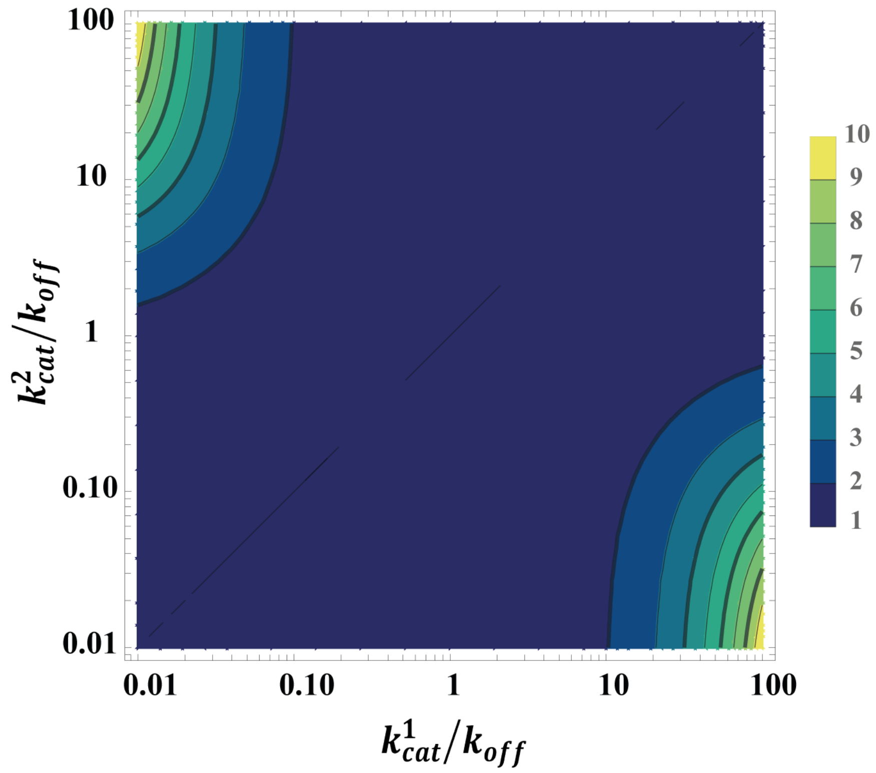


**Figure S3:** 2D color map of the error factor $\frac{\langle W_{ES} \rangle}{\langle W_{ES}^{cat} \rangle}$ calcuted in the space of the normalized catalytic rates $\frac{k_{cat}^{1}}{k_{off}}$ and $\frac{k_{cat}^{2}}{k_{off}}$. The calculations were performed for a splitting probability of $p = 0.9$.
